# Sea urchin consumption of kelp controlled by the density of drift algae

**DOI:** 10.1101/2025.11.09.687392

**Authors:** Zachary Randell, Casey Sheridan, Jessica Frey, Mark Carr, Mark Novak

## Abstract

Rocky reef kelp forests can exhibit abrupt shifts between forested and barren states where unchecked sea urchin grazing inhibits kelp recovery. Forefront to urchin barren formation is a behavioral switch where sedentary urchins become active and graze upon attached kelp. This is hypothesized to occur when the availability of drift (i.e., detached kelp on the surface of the reef), which urchins are believed to prefer over intact and attached kelp, declines. Although observed repeatedly, the density-dependent nature of this apparent resource preference has not been characterized quantitatively. We used a series of subtidal functional response experiments in central California, USA, to quantify how consumption rates by *Strongylocentrotus purpuratus* (purple urchins) on *Macrocystis pyrifera* (giant kelp) and drift varied with their relative availability and over time as urchins became satiated. Our experiments indicate that the preference for drift over kelp is strong when drift and kelp are at equal abundance, that the abundance of drift has a strong effect on kelp consumption but not vice versa, and that the preferences for drift is indeed density dependent with the qualitative switch to a greater preference for kelp occurring when the availability of drift declines below approximately 1 *g* of drift for every 60 *g* of kelp. Our analyses suggest further that maintaining at least equal amounts of drift and kelp is sufficient to all but preclude urchins from grazing on live kelp. We recommend drift be used as an indicator of reef health, and that our thresholds be considered to guide drift subsidies to preserve and restore kelp forests.

## Introduction

Managing ecological communities that exhibit alternative stable states requires an understanding of what drives the transitions between states and of the feedback processes that stabilize within-state dynamics to inhibit the recovery of desirable states (Thomas, 1981; Scheffer *et al*., 2001; Schröder *et al*., 2005; van de Leemput *et al*., 2016). Temperate rocky reefs are renowned for their capacity to exhibit multiple stable states, including shifts between diverse and productive communities of macroalgae (i.e., kelp forests) and relatively depauperate states dominated by encrusting coralline or turf algae. In cases where the shift is associated with and maintained by urchin grazing, this deforested and generally less desirable state is known as an urchin barren (Laurence, 1975; Mann, 1977; Filbee-Dexter & Scheibling, 2014; Randell *et al*., 2022). These state shifts are proximately caused by disturbances such as large wave events and are compounded by stressors like nutrient limitation and elevated temperatures, the combinations of which detach kelp, reduce its growth, and inhibit the recruitment of younger stages (Dayton *et al*., 1992; Steneck *et al*., 2002; Reed *et al*., 2011; Boada *et al*., 2017).

In kelp forests dominated by giant kelp, *Macrocystis pyrifera*, most standing biomass is not consumed directly by herbivores (Mann, 1973; Schiel & Foster, 2015). Instead, a significant fraction senesces or is dislodged to become drift algae (Krumhansl & Scheibling, 2012a). This drift is generally retained within kelp-forest patches, providing a key resource subsidy for benthic consumers including urchins. Crucially, drift availability is believed to mediate urchin grazing behavior. Despite similarly high overall urchin densities in healthy forests and barren grounds, urchins remain sedentary in crevices and sheltered habitats when drift is sufficiently abundant, passively consuming drift rather than actively grazing on attached kelp (Harrold & Reed, 1985; Randell *et al*., 2022). However, when drift availability declines, urchins exhibit a marked behavioral shift, emerging from shelter to actively forage on attached kelp, potentially precipitating a transition toward an urchin barren (Ebeling *et al*., 1985; Smith & Tinker, 2022). The behavioral shift is often particularly pronounced in locations where urchin consumers are absent or at low abundance (Filbee-Dexter & Scheibling, 2014; Eisaguirre *et al*., 2020; Ortiz-Villa *et al*., 2025; Pessarrodona *et al*., 2019).

Effects of drift availability on urchin movement and grazing have been demonstrated by prior experiments and long-term monitoring. For example, the laboratory experiments of Kriegisch *et al*. (2019) showed that urchins exhibit directional movement towards drift and to subsequently decrease their movement rate when in the presence of drift. Similarly, Vanderklift & Kendrick (2005) demonstrated that drift tends to accumulate alongside sedentary urchins and that urchins consume more drift than attached kelp, such that urchins had minimal impact on the abundance of attached kelp. Via long-term monitoring, drift has also been shown to modify kelp biomass, annual variation in kelp dynamics, and the propensity for the system to exhibit a switch towards urchin barrens (Rennick *et al*., 2022). However, despite a general recognition that urchins will consume drift more readily than attached kelp, the density-dependent nature of urchins’ resource preference — defined as the disproportionate consumption of a resource relative to its availability (Chesson, 1978) — remains uncharacterized. Specifically, it is unknown how the availability of drift and kelp interact to influence the threshold at which urchins switch (sensu Murdoch, 1969) from passive drift consumption to active kelp grazing.

To address this gap, we performed a series of subtidal caging experiments in central California that systematically varied the abundance of drift and attached kelp and quantified urchin consumption rates over time. Employing a novel model-fitting analysis incorporating time-dependent resource restocking and non-equilibrial urchin satiation, we quantitatively characterized urchin resource preferences to demonstrate that urchins exhibit a strong baseline preference for drift over attached kelp when the two resources are equally abundant. Our analysis revealed that this preference is highly density-dependent: urchins disproportionately consume drift over kelp when drift is abundant, and shift to grazing attached kelp when drift availability declines below approximately 1 *g* of drift per 60 *g* of attached kelp. Our results further indicate that kelp consumption rates strongly depend on drift availability but that drift consumption remains largely independent of kelp abundance over a large range of abundances. These findings provide an empirically informed basis for better modeling consumer-resource dynamics and offer quantitative thresholds to guide the use of drift subsidy and retention strategies for preserving and restoring kelp forests.

## Methods

### Field experiments

The experiments were conducted near the Hopkins Marine Station in Monterey, California, USA, over the summer of 2019. We affixed an array of 20 cages onto the seafloor at depths of 4-6 *m* in the center of a large sandy area between two rocky reefs (36^°^37^′^12^′′^N, 121^°^54^′^07^*′′*^W). The cages were comprised of a single 7.62 *m* length of steel rebar bent into a 1 x 1 x 0.4 *m* frame (Fig. 1*a,b*). Two layers of netting and screening enveloped the cages and prevented transit of particulate matter greater than 2 *mm*^2^ into or out of the cages. Eight paving stones were placed into four stacks of two within each cage to form a “plus-sign” shaped crevice in which urchins could hide (Fig. 1*c,d*). See Figs. S1 and S2 and the *Supporting Information* for details.

**Figure 1:**
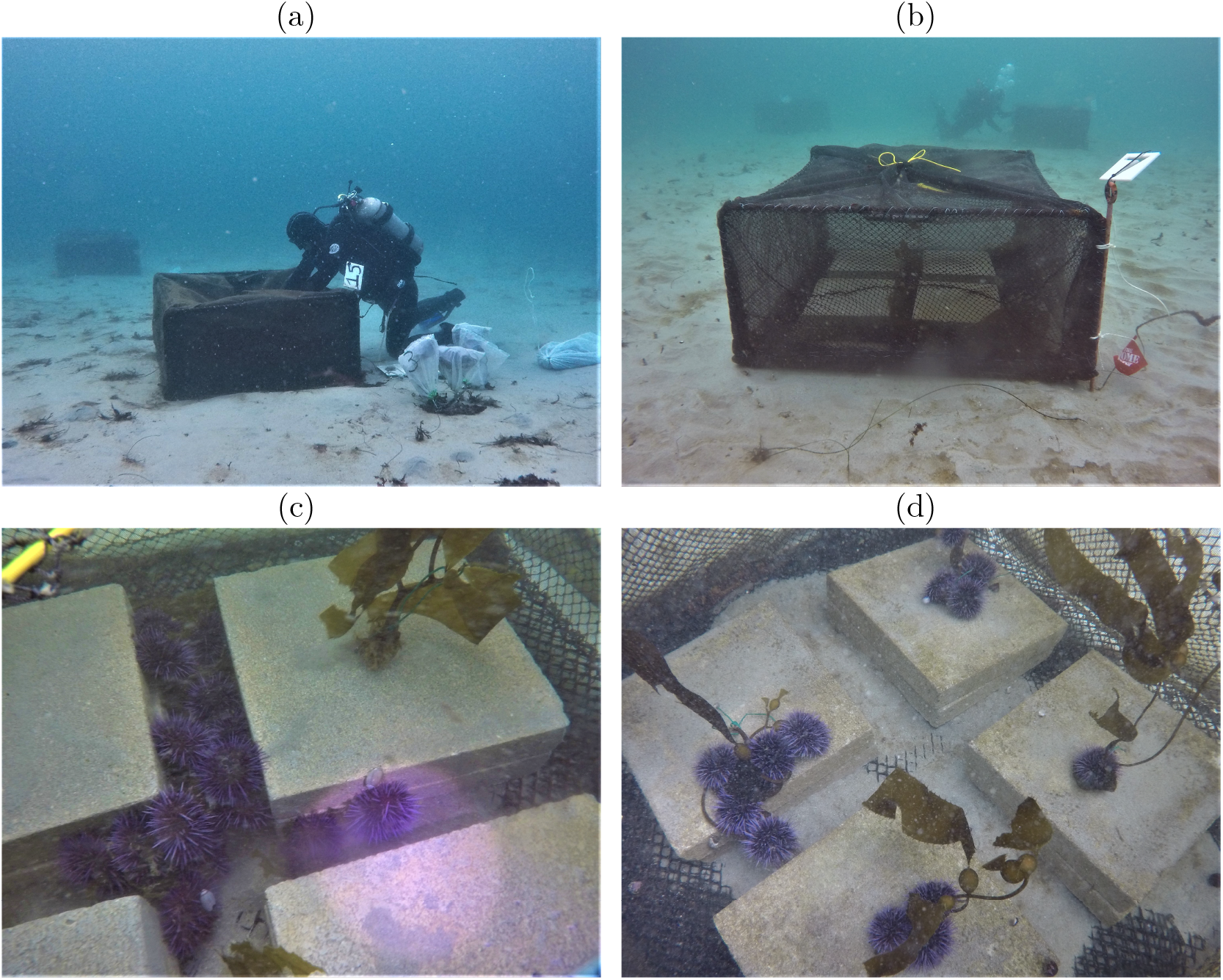
(*a*) A diver accesses the interior of a cage through a window in both the screening and netting. (*b*) A cage with the four stacks of two paving stones. (*c*) Urchins consuming drift in the “plus sign” of the paving stones. (*d*) A treatment with urchins actively grazing kelp atop the paving stones.

In each cage we placed either 10 or 20 purple urchins of 4-5 *cm* diameter, collected by hand from barrens to the northwest of Hopkins. Urchins were allowed to habituate to their cage for 24 *h* before the onset of a trial. We provided abundant drift for the urchins to consume during this period and removed all remaining drift prior to initiating treatments. This served to standardize gut fullness (Griffen, 2021) and induce the sedentary behavior reflective of the forested state. We then added drift and kelp to each cage, targeting a factorial combination of 9 levels of drift (see below) and three levels of kelp: no kelp (i.e., drift only), a “low” range of 10-130 *g* wet weight consisting of a single kelp individual, and a “high” range of 100-330 *g* wet weight consisting of two or four individuals (average weight per individual: 53.05 *g*, std dev: 25.05 *g*; see Table S1 and Figs. S3 and S4 for context and details). Drift was comprised of adult blades of Giant Kelp (*Macrocystis pyrifera*) sporophytes that were clipped, weighed, and gently rolled into a goodie bag for transport to the cages within 3 *h* of collection. Kelp were young *M. pyrifera* sporophytes (*<*1 *m* length) that were carefully removed from the natural substrate with their holdfast intact. Kelp holdfasts were fixed to the top of the paving stones using wire (Fig. 1*d*), replicating their typical location atop rocky reefs.

The above design was used in a total of eight replicate trials (Fig. S5 and Table S1). For the first four of these trials (*Sequence 1*), we varied initial drift abundance across a range of no drift (i.e., “low” or “high” kelp only) or 10-330 *g* wet weight and quantified remaining kelp and drift biomass after approximately 44, 89, and 134 *h*. For the remaining four trials (*Sequence 2*), we varied initial drift abundance across a range of no drift or 10-110 *g* wet weight and quantified remaining kelp and drift biomass after approximately 18 and 36 *h*. The range of drift abundances we used is representative of the densities seen on natural reefs (Harrold & Reed, 1985; Wernberg *et al*., 2006; Smale *et al*., 2022).

Kelp and drift were restocked between time periods to within 10% of their original trial abundance. Urchins were not replaced between time periods to track cumulative consumption through time. However, we did use new urchins from the barrens and re-randomized cage-specific treatment assignments for each trial. Logistical field errors for 5 observations in *Sequence 1* and for 1 observation in *Sequence 2* caused us to remove their respective treatments (i.e., cages) from our analysis. Therefore, in total, our data for *Sequence 1* consisted of 71 cages followed over three time periods (213 data points) and for *Sequence 2* consisted of 75 cages followed over two time periods (150 data points).

### Model fitting

We fit a suite of eight competing models to our experimental data to quantify how urchin consumption rates on kelp and drift varied as a function of their abundances over the time course of the trials. The models accommodated the potential for several key features we expected to see, including preferences that changed with the relative abundance of the two resources, grazing rates that declined as urchin movement rates declined at high abundances of their preferred resource, and grazing rates that declined as urchins became satiated over time. The eight models differed from each other in (*i*) their assumed preference function (either a logistic model or a reduced version of the preference function underlying the functional response model of van Leeuwen *et al*. (2013)), (*ii*) the resource-dependence of urchin movement rates (either dependent or independent of resource abundances), and (*iii*) the consideration of the gut evactuation process (either explicit or implicit).

#### Base model

Our models described the treatment-specific rates of change in the observed abundances of drift *S* and kelp *A*, as well as the average urchin’s unobserved gut fullness *F* (i.e., a latent variable), through a system of differential equations:

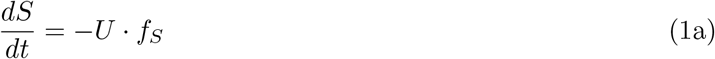

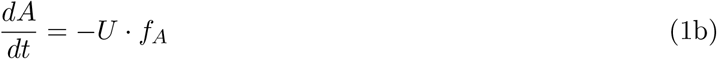

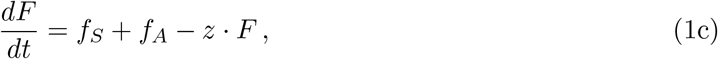

where *U* is the number of urchins in a cage, *f*_*S*_ and *f*_*A*_ are the average urchin’s consumption rates on drift and kelp respectively, and *z* is the gut evacuation rate (explicitly modeled). Consumption rates (i.e., the average urchin’s functional response on each resource) were modeled as

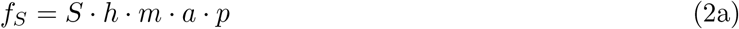

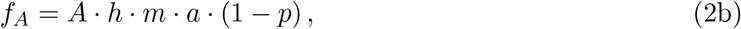

where *a* is the urchin’s intrinsic resource search rate, gut fullness controlled the urchin’s hunger level as *h* := *h*(*F*) = exp[−*v*· *F*] for which *v* represents the gut satiation sensitivity, and the urchin’s realized movement rate *m* was determined by

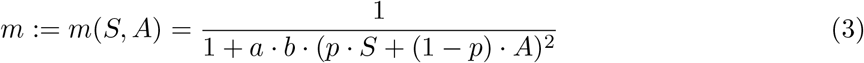

for which *b* represents a movement suppression rate and *p* the preference for drift. The urchin’s preference for drift versus live kelp was either described by either a logistic preference function that underlies the resource-switching functional response model of Koen-Alonso (2007, which he attributes to Peter Yodzis) or the preference function underlying the functional response model of van Leeuwen *et al*. (2013) reduced to two resources. These functions are mathematically equivalent to

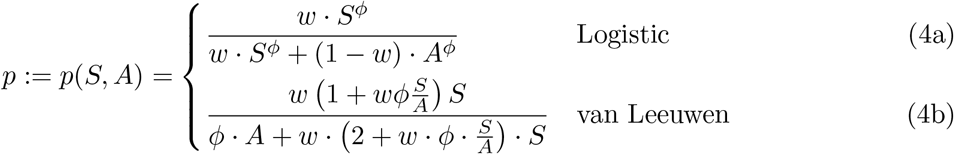

(see *Supplementary Information*).

Our base model thus assumed that urchins exhibit an overall resource search rate of *a* reflecting their average movement velocity and per capita grazing rate when fully hungry at low resource abundances; that grazing rates could decline over time depending on the average urchin’s hunger level *h* as determined by its gut fullness *F* and gut satiation sensitivity *v*, with gut fullness determined by consumption and a gut evacuation rate proportional to fullness; that movement and resulting search rates could decline at high densities of the preferred resource depending on the movement suppression rate *b* (described in a phenomenological manner similar to the type IV Monod-Haldane model (Andrews, 1968) with satiation itself being represeted by *h* and *F*, see *Supplementary Information*); and that the average urchin’s preference *p* for encountered drift versus its preference (1 − *p*) for encountered live kelp was determined by the ratio of drift to kelp abundances. That is, while *p* was set to 1 for the drift-only controls and to 0 for the kelp-only controls, the urchin’s preference for drift could otherwise change from a baseline preference of *w* when drift and kelp were equally abundant by disproportionately increasing for higher ratios of available drift and kelp at a switching sensitivity *ϕ*. Values of *ϕ >* 1 would affect disproportionately higher preferences for drift as its relative abundance to kelp increases (i.e., positive switching), while values of *ϕ <* 1 would affect disproportionately lower preferences for drift as its relative abundance increases (i.e., negative switching). We note that our modeling of gut fullness as a dynamic variable permitted us to avoid the assumption inherent in functional response models that the urchins’ behavioral states of searching, handling, and digesting were at an equilibrium (Jeschke *et al*., 2002; Novak *et al*., 2025) though this was not so for the phenomenological representation of movement suppression.

#### Specific models

Half of our eight models assumed the logistic preference function. The other half instead assumed the simplified van Leeuwen preference function. For each of these two preference functions, we also fit a model in which urchin movement rates were assumed to be constant and independent of resource abundances (setting *m* = 1, equivalent to *b* = 0), a model in which the gut evacuation process was considered to be implicitly represented by the urchin’s hunger level (setting *z* = 0, interpreting the effective gut fullness *v* · *F* as the net effect of gut filling and evacuation), and a model that combined these assumptions (setting both *m* = 1 and *z* = 0).

#### Model fitting and comparison

We fit these models to our time series of initial, measured, and restocked kelp and drift abundances using a Bayesian approach to obtain probabilistic estimates of the parameters and predicted consequences. Parameter priors were specified to be uninformative in determining the urchins’ preferences for drift versus kelp, and weakly informative in so far as determining the order-of-magnitude difference in the scales of their search rate, gut clearance rate, gut satiation sensitivity, and movement supression rate. The performance of the models was assessed on the basis of their leave-one-out estimates of the expected log pointwise predictive density (Vehtari *et al*., 2025; Sivula *et al*., 2025), with model weights determined by the pseudo-BMA+ method (Yao *et al*., 2018).

## Results

### Field experiments

Urchins in treatments with abundant drift (≳ 100 *g*) settled into the base of the cages, either in the “plus-shaped” crevice between the paving stones or between their perimeter and the cage siding (Fig. 1*c*). Active urchin movement was extremely rare in these treatments. In contrast, when drift was scarce, urchins actively moved over all interior surfaces of the cage, including atop the paving stones. We observed these urchins grazing on kelp stipes just above the holdfast (which sometimes detached the frond and rendered it unreachable when it floated to the top of the cage), climbing up onto erect fronds to graze, and pinning fronds to the substrate to graze (Fig. 1*d*).

In the trials of *Sequence 1*, the average amount of drift that was consumed increased as a function of initial drift abundance and decreased across the time periods. For initial drift near 100 *g* or less, drift consumption reached maxima of approximately 3.5, 2.75, and 1.25 *g* urchin^−1^ 48 *h*^−1^ in periods 1, 2, and 3, respectively (Fig. 2 row 1). For initial drift above 100 g, drift consumption declined in periods 1 and 3, but in all periods was independent of the abundance of kelp. Kelp consumption also decreased across the time periods and was likewise dependent upon the availability of drift (Fig. 2 row 2). Kelp consumption was independent of kelp abundance (Fig. 2 row 3). Consumption rates in the shorter-duration trials of *Sequence 2* exhibited similar patterns across the narrower range of provided drift abundances (Figs. S7 and S8).

**Figure 2:**
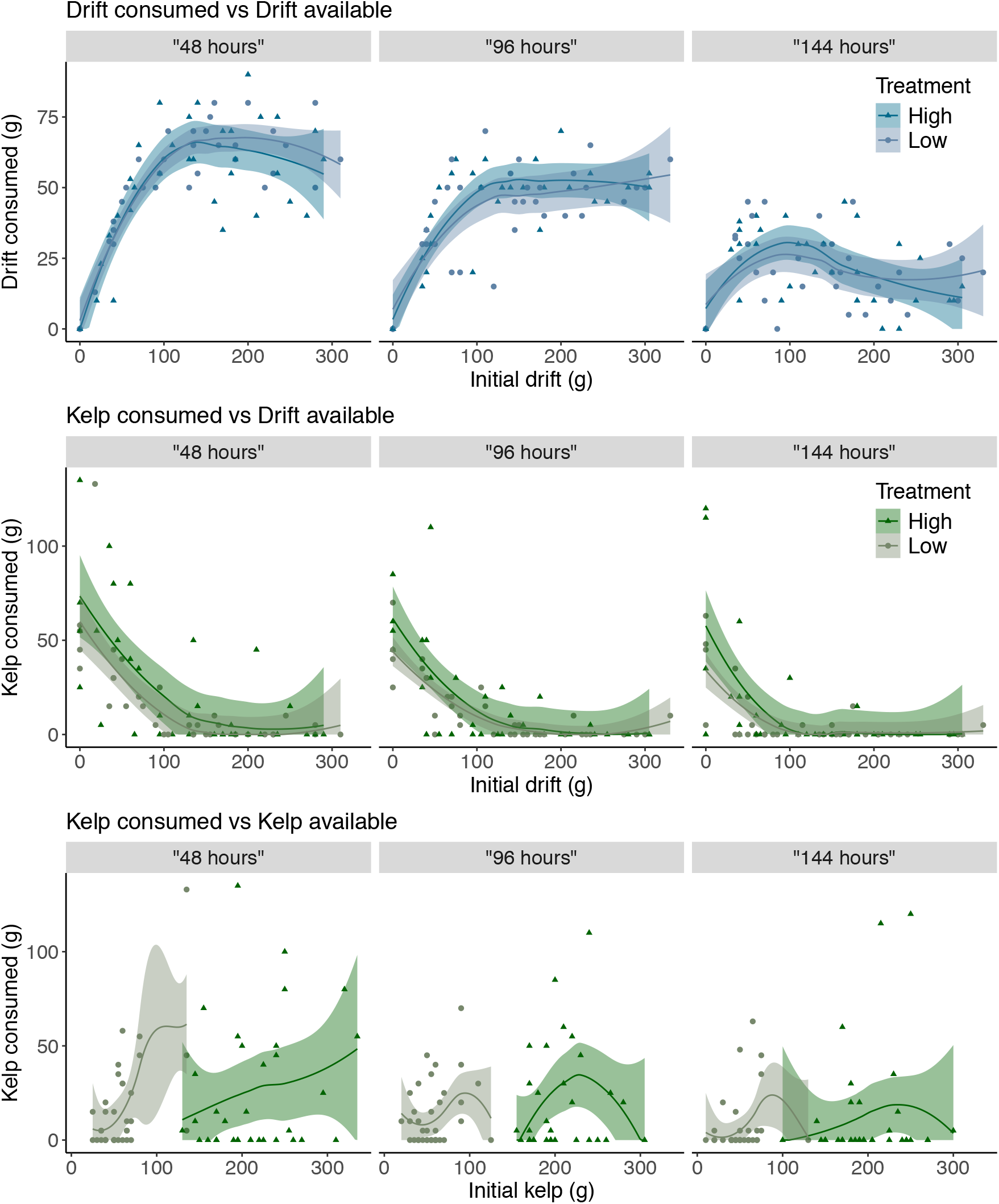
The relationship between initial drift available and drift consumed (row 1), between initial drift available and kelp consumed (row 2), and between initial kelp available and kelp consumed (row 3) across the three time periods of *Sequence 1* distinguishing “low” (one kelp plant) and “high” (four kelp plants) treatments. Ribbons reflect loess-smoothed means (span = 1) and 95% confidence internals. Rows 1 and 2 also depict the 24 control treatments of 1 or 4 kelp individuals to which no drift was added; these were not included in the loess smooths and model fitting. See Fig. S8 for *Sequence 2* data.

### Model fitting

All four models that allowed for a suppression of urchin movement at high abundances of their preferred resource performed similarly well in replicating the patterns of kelp and drift consumption that we observed across the resource abundances and time periods of the experiments (model weights: 0.16-0.39); those that did not allow for the suppression of urchin movement received no support (Table S2, Figs. S11-S18, parameter estimates given in Table S3). We therefore used model-averaging based on model weight to incorporate the structural uncertainty among the top four models into our subsequent inferences.

At illustrative initial abundances of 50 *g* and 250 *g* for “low” and “high” kelp treatments respectively, the predicted drift consumption of all four models (and thus their average) exhibited an overall decrease across time periods, a saturating increase at low initial drift abundances that reached a maximum near 100 *g* of initial drift within each time period, and a relative insensitivity to variation in kelp abundance (Fig. 3*a-c*). Although model-predicted kelp consumption was more sensitive to kelp abundance at low drift abundances than was observed in the experiments, the models well-described its dependence on drift abundances at low kelp abundances as well as its overall decrease across the time periods (Fig. 3*d-f*). Underlying these dynamics in consumption rates were inferred changes in the urchins’ resource preferences (see next) and their gut fullness. The latter consistently increased across the time periods, was generally higher at high than at low kelp abundances when drift abundance was low, and showed a unimodal relationship with drift abundance (especially at low kelp abundance; Fig. 3*g-i*).

**Figure 3:**
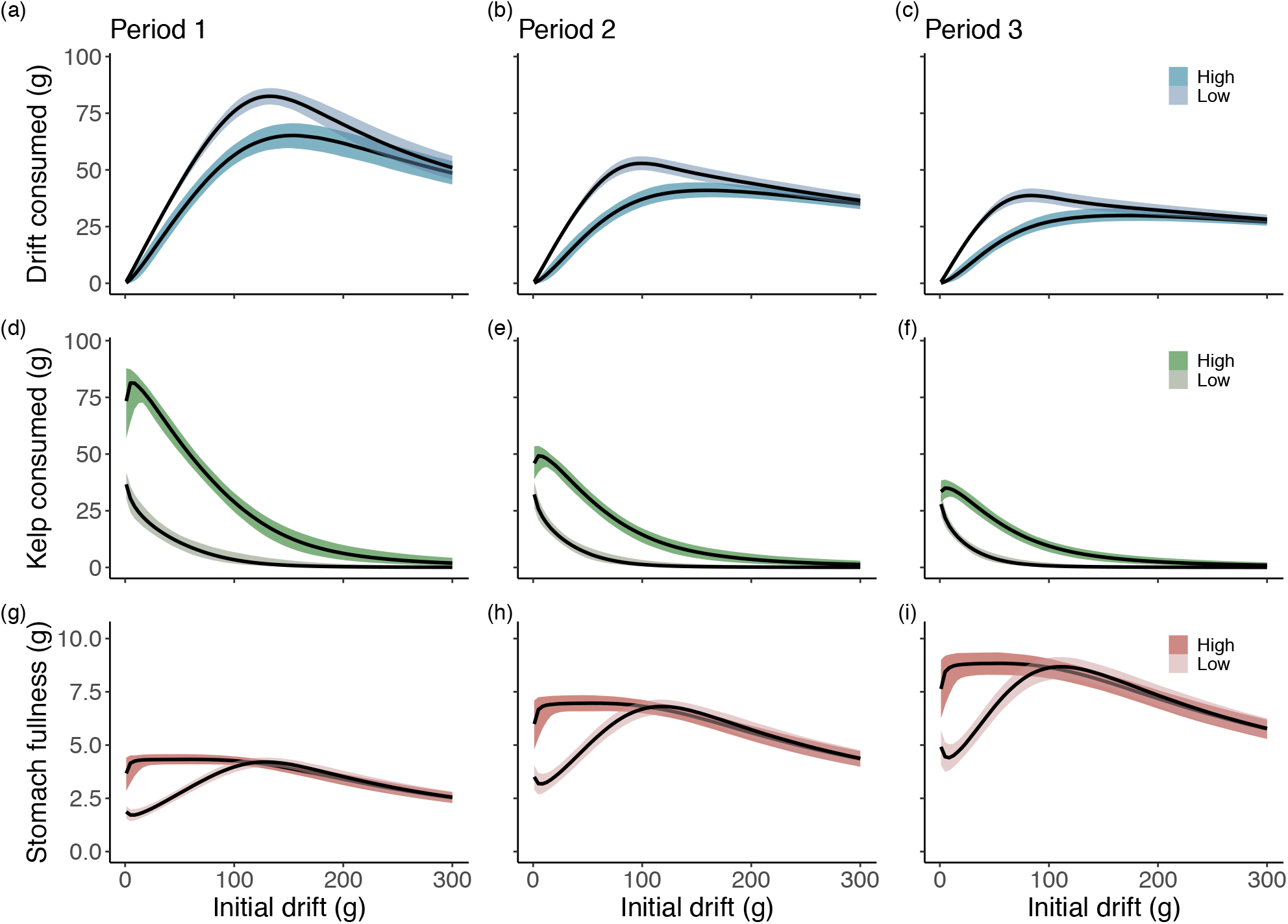
The model-averaged median predicted consumption of drift and kelp and corresponding cumulative gut fullness as a function of initial drift abundance under illustrative “low” (50 *g*, lighter color) and “high” (250 *g* darker color) initial kelp abundances. Ribbons reflect 95% credible intervals. Period 1: 44 *h* (*a,d,g*); period 2: 89 *h* (*b,e,h*); period 3: 134 *h* (*c,f,i*). See Figs. S15-S18 for model-specific predictions.

On the basis of the weight-averaged model, urchins were inferred to exhibit a very strong baseline preference for drift over kelp, consuming drift and kelp at a median relative rate of 22.2 to 1 (*p* = 0.96; 95% credible interval: 0.91-0.98) when drift and kelp were at equal abundance (Fig. 4, Table S4). Below this relative resource abundance their preference became strongly dependent on the relative abundance of the two resources, exhibiting a qualitative switch to a stronger preference for kelp below a median abundance ratio of 1 *g* of drift to 56 *g* kelp (95% credible interval: 14-5624 *g*). Each of the models that allowed for movement suppression provided similar inferences to the these model-averaged inferences regardless of the assumed preference function and how gut evacuation was described (Table S4), although the van Leeuwen preference function did associate a higher probability to urchins retaining a stronger preference for drift even at very low drift to kelp ratios (Figs. S19-S22).

**Figure 4:**
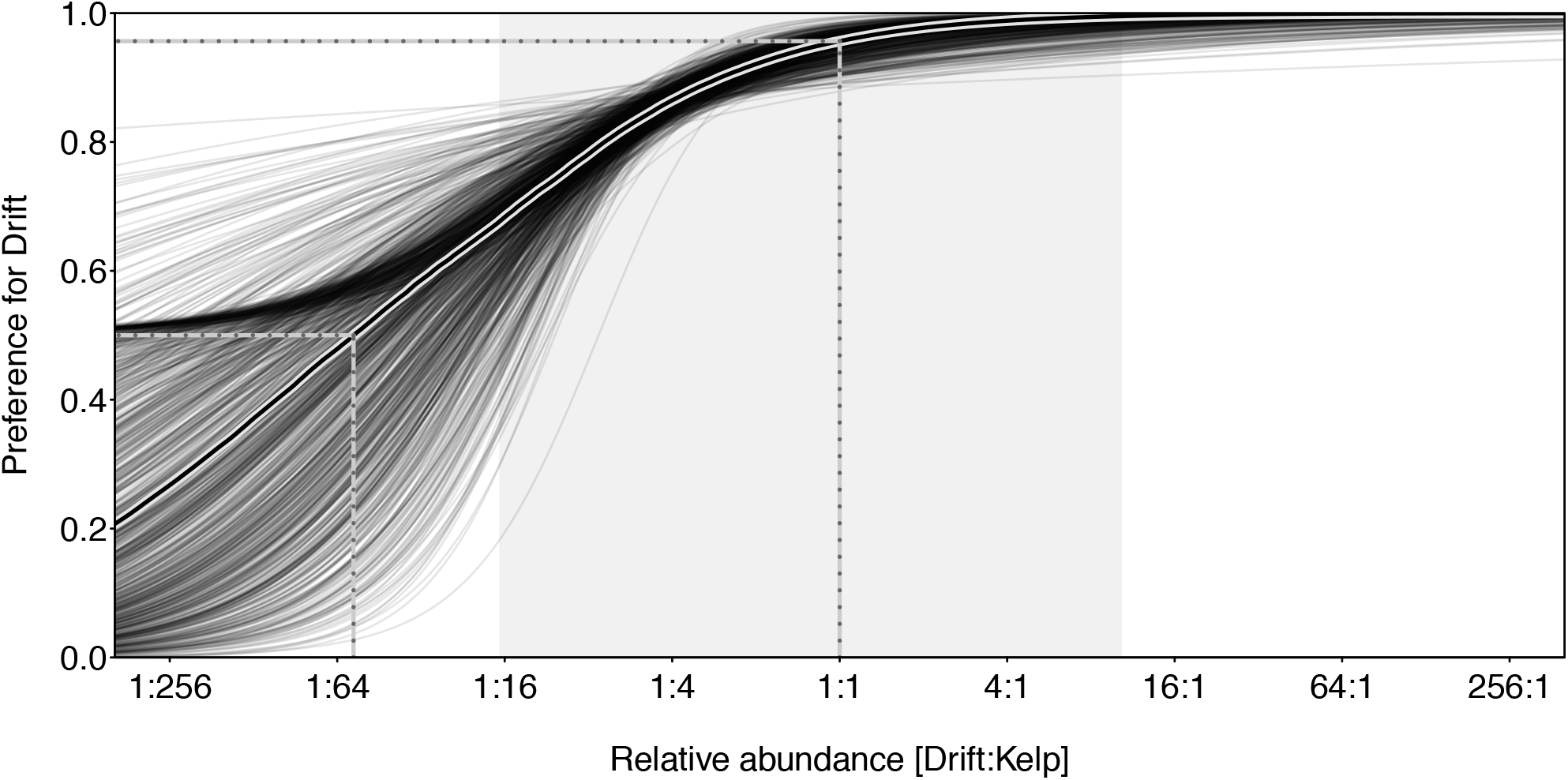
Model-averaged predictions of urchin resource preference for drift as a function of the relative abundance of drift and kelp biomass. Each thin black line reflects a posterior draw, with the thick central line indicating the median preference. The shaded gray region indicates the range of relative abundances in the experiment. The central/upper dotted lines mark the estimated preference for drift of corresponding to equal (1:1) resource abundances. The left/lower dotted lines mark the point of equal preference (*p* = 1 − *p* = 0.5). See Figs. S19-S22 for model-specific predictions.

## Discussion

Our experiments demonstrate that the abundance of drift controls the rate at which urchins consume kelp. When even modest quantities (≳ 60 *g*) of drift were available, urchins overwhelmingly consumed it rather than kelp (Fig. 2). Our models well-reproduced the temporal progression of the experimental patterns (Fig. 3), revealing that urchins preferred drift with near exclusivity when the two resources were equally abundant and did not exhibit an equal preference for them until drift biomass fell below about a sixtieth the biomass of kelp (Fig. 4). Our work therefore corroborates prior work in demonstrating that drift acts as a powerful regulator of urchin behavior capable of insulating kelp from urchin grazing, but also provides quantitative targets for drift abundances with which to inform kelp forest management and restoration.

Both our models and our direct observations of the experiments suggest that the urchins’ strong preference for drift was mediated by their rates of movement. Urchins with all but the lowest levels of drift quickly wedged themselves into the “plus-sign” and outer sides of the paving stones, largely remaining immobile while consuming drift. In contrast, drift-deprived urchins, which at first also behaved this way, subsequently left shelter, climbed the cages vertical sides, and actively grazed the live kelp atop the paver stones in a manner characteristic of the active grazing associated with barren formation. Their reduction in movement and the coincident preference for drift when drift was available was unlikely due to differences in the composition or nutrient profile of the drift and kelp given that the two were the same species in our experiment (albeit at different life-stages and states of senescence/decay). Rather, we believe it reflects the urchins’ behavioral predilection to make use of habitat features that provide perceived protection against predators (Tegner & Levin, 1983; Nichols *et al*., 2015; Smith *et al*., 2021). Active wandering and barren formation are thereby more a consequence of hunger than a shift in dietary preference for kelp versus drift per se. Thus, although high amounts of drift (≳ 100 *g*) were the most effective at reducing the urchins rates of movement (Fig. 2), even modest rates of detrital influx are likely to be sufficient to satisfy their low energetic demands and preclude active movement (Kriegisch *et al*., 2019; Vanderklift & Kendrick, 2005).

The repeated restocking of resources in our experiments was important for disentangling the effect of resource abundances on urchins’ movement, preference, and satiation over time. This necessitated the use of a more complex, non-traditional approach to functional response model fitting. That is, although ordinary differential equations like eqns. (1a-b) have been used to model resource depletion and influx before (e.g., Novak, 2010; Rosenbaum & Rall, 2018), our study’s explicit consideration of the restocking events that dynamically perturbed what would otherwise have been uninterrupted declines in resources was novel. Doing so not only added empirical insight into the temporal progression of the urchins behavior, but — with the exception of the gut evacuation rate — also increased the information available for estimating the model’s parameters (see also Capitán & Alonso, 2025). Our use of a latent variable for gut fullness likewise expanded upon existing functional response model-fitting approaches regarding consumer satiation. Since the pioneering models of Ivlev (1961) and extending to the present, satiation has been assumed to be at steady state — and thus constant with respect to time for any given resource abundance — in the derivation of functional response models (see Jeschke *et al*., 2002; Novak *et al*., 2025). At the expense of additional parameters whose values we were able to estimate by virtue of resource restocking, our model avoided this assumption by considering hunger level to be a function of cumulative rather than instantaneous ingestion.

Our threshold estimates of urchin behavior should nevertheless be interpreted in the context of our experiments, rather than being seen as intrinsic to urchins universally. This is because our experiments sought to standardize physical and biological conditions within 1 x 1 *m in situ* cages — including substrate configuration, refuge availability, water motion, urchin density, and predator access — in order to isolate the effects of varying drift and live kelp densities on urchin consumption. Therefore, although the ranges of kelp and drift abundances we supplied were representative of their densities on natural reefs (Fig. S3; Harrold & Reed, 1985; Wernberg *et al*., 2006; Smale *et al*., 2022) and the crevice-like refuge we provided allowed urchins to exhibit their natural behavior, the expression of this transition on natural reefs will likely vary across differing physical, ecological, and physiological contexts.

For example, substrate complexity can affect hydrodynamic forces, drift retention, and urchins’ access to drift and attached kelp (Laur *et al*., 1986; Vadas *et al*., 1986; Randell *et al*., 2022), and can alter urchin movement rates directly, with urchins reducing movement and increasing their reliance on structural refuges as swell increases (Lauzon-Guay & Scheibling, 2007; Parnell *et al*.,2017). Likewise, urchin density itself can also affect urchin movement and responsiveness to resource subsidies (Parnell *et al*., 2017), temperature can influence urchin metabolic rates and responsiveness to kelp-derived chemical cues (Mann *et al*., 1984), and predators and predator-associated cues can suppress movement and grazing activity (Harrold & Reed, 1985; Vadas *et al*., 1986; Pessarrodona *et al*., 2019). Notably, the urchins we used were sourced only from an urchin barren; because kelp-forest urchins are typically better fed, healthier, and frequently gravid (Dolinar & Edwards, 2021; Grime *et al*., 2023), they may require less drift to remain sedentary and may delay active grazing even as drift becomes scarce. Repeating our experiments with kelp-forest urchins would help determine how urchin condition modifies the drift availability needed to suppress active grazing. Extending such manipulations to also encompass environmental gradients and larger spatial extents would clarify how urchins’ preference for drift varies among reef contexts and would greatly aid in the translation of our cage-scale results to the scales more immediately relevant to management and restoration.

Caveats aside, our study adds to a growing literature identifying drift as a key determinant of urchin behavior and kelp forest persistence. Drift availability may therefore serve as a measurable indicator of a kelp forest’s capacity to buffer disturbances and preclude shifts toward active urchin grazing, particularly because variables closely linked to the positive and negative feedbacks that determine the stability of alternative states may often be used to forewarn of critical transitions (Scheffer *et al*., 2009; Nijp *et al*., 2019; Kéfi *et al*., 2014, but see Dakos *et al*. 2015). Fine-scale estimates of drift availability within kelp forests will likely require diver-based surveys given the difficulty of distinguishing detached drift from attached macrophytes in benthic imagery, but imaging platforms such as remotely operated vehicles will likely be useful for assessing drift availability across larger spatial scales beneath reefs or along reef margins where drift often accumulates (Britton-Simmons *et al*., 2012; Filbee-Dexter & Scheibling, 2016; Krumhansl & Scheibling, 2012b).

Our study does suggest the possibility of subsidizing drift to mitigate urchin overgrazing, or temporarily divert urchins away from outplanted kelp, in the contexts of kelp forest conservation and restoration (Watanuki *et al*., 2010; Morris *et al*., 2020; Gleason *et al*., 2021). Subsidizing drift is not without significant logistical, ecological, and management complexities, but creative solutions may exist. Along the northeast Pacific coast, certain areas with giant kelp may support harvesting to create drift, though this would not be an option for areas with bull kelp (*Nereocystis luetkeana*) for which harvesting from the upper canopy removes reproductive capacity. Similarly, removing drift directly from one reef to supply another may lead to unintended consequences for source locations. Kelp aquaculture could provide an abundant and seasonally consistent supply of drift, while other forms of aquaculture — such as shellfish farms and integrated multi-trophic systems — often produce excess algal growth that does not have immediate use (Grebe *et al*., 2019; Green-Gavrielidis *et al*., 2023; Ulaski & Konar, 2024). Harvesting beach wrack may be another option in limited cases. Additionally, because urchins are omnivorous, non-macrophyte resources, such as fish byproducts (Sivertsen *et al*., 2008) offer additional alternatives.

While logistical limitations exist, subsidizing drift ought to be considered an option, particularly along seascapes with degraded top-down urchin control. Given spatiotemporal limitations in the extent to which drift subsidies could realistically be applied, they should not be considered a large-scale solution for restoring urchin barrens to kelp forests. Rather, we posit they could be a short-term, targeted strategy to modify urchin behavior at critical junctures in conservation and restoration processes. For example, the success of kelp restoration could be increased by diverting urchins to subsidized drift, especially when paired with urchin removals (e.g., McHugh *et al*., 2025). Such a three-pronged approach — urchin removal, kelp restoration, and drift provisioning — could buy time for kelp enhancements to establish self-replenishing densities, including the innate production of drift (White *et al*., 2026). Drift subsidies could also be used more defensively to buffer remnant kelp patches which may be disproportionately important for future recovery given the metapopulation dynamics that characterize kelp forests (Castorani *et al*., 2015, 2017; Ricart *et al*., 2025). In this context, limited “firewalls” of drift could help stave off grazing fronts, preserve local spore sources, and reduce the likelihood that remnant kelp patches transition into persistent barrens, particularly when conducted alongside urchin removals. These approaches may be particularly relevant along the northeast Pacific coast where key urchin predators, such as sea otters, sunflower stars, sheephead, and spiny lobster, have either been extirpated, devastated by disease or reduced by fishing, eroding the ecosystem’s capacity to curb urchin grazing (Hamilton & Caselle, 2015; Burt *et al*., 2018; Langendorf *et al*., 2025). Indeed, the importance of resource-driven behaviors documented here has likely become magnified in many ecosystems where historically strong top-down controls have waned (Estes *et al*., 2011).

## Acknowledgments

We are grateful to Heather Fulton-Bennett, Bryant Gutierrez, Hannah Knotter, Anya Morrill, Benji Nguyen, Samuel Tse, and Hailey Whitehead for their efforts in the field, and to members of the Stan Project team for their coding assistance. We are also grateful to Hopkins Marine Station for hosting our experiment and granting us access to their much-appreciated warm showers, and to the Partnerships for Interdisciplinary Studies of Coastal Oceans (PISCO) and the Raimondi-Carr Lab for their support. We are also grateful to the U.S. Geological Survey Western Ecological Research Center, Santa Cruz field station, for providing access to a vessel. Urchins were collected per California Department Fish and Wildlife Scientific Collecting Permit S-190860002-19086-001. ZR was supported by a NSF Graduate Research Fellowship (2016227734), PISCO, and the Dr. Earl H. & Ethel M. Myers Oceanographic & Marine Biology Trust. MN was supported by a David and Lucile Packard Foundation grant to PISCO, the NSF (DEB-2129759), and the Oregon Ocean Science Trust. This is PISCO publication #554.

