## Supplementary Material for "Sea urchin consumption of kelp controlled by the density of drift algae"

**References** **31**

### S1 Experiment details

#### Cage and treatment design

Each subtidal cage was made of a single 20 $ft.$ length of rebar manually bent into a $1 \times 1 \times 0.4\ m$ frame (Fig. S1$a$) using a vice. Hard plastic Vexar comprised the floor of the cage. Two bungee cords provided flexible support to the two upper sides of the cage not consisting of rebar. A single zinc anode was attached to the rebar frame. Knotted nylon netting (mesh size 2 $cm$) was stitched around the entirety of the cage via abrasion resistant braided fishing line (Fig. S1$b$). The cage was then completely enveloped in fiberglass window screening which was stitched onto the rebar frame like the nylon netting (Fig. S1$c,e$). The "double-hull" of netting and screening prevented particulate exchange ($> 2\ mm$) in or out of the cage. Although it likely decreased light availability, the effect on kelp growth was likely negligible given the short temporal scale of our experimental trials. Although water motion was likewise attenuated, the effect was not enough to prevent drift from visibly shifting about on the cage bottom with the surge. Both screening and netting "doors" overlapped together on the top center of the cage, allowing divers to quickly and easily access the entirety of the cage interior (Fig. S1$a$). Aluminum grommets were used to allow the screening to be cinched shut with paracord (Fig. S1$c,d,f,g$). This ease of access was essential for meticulously collecting all fragments of drift (including from underneath urchins) during the experimental trials. A single 1 $m$ Earth anchor was augured into the sand and lashed to the rebar frame along one corner of the cage. Additional cage support was provided by the four stacks of two paving stones (each 20 $lbs$) inside the cage. The cages were spaced 5 $m$ apart in a rectangular grid. 1 $m$ lengths of rebar with labeled attachment points were placed next to the cage array to organize treatment application and removal.

The 30 - 300 $g$ range of drift was selected based on preliminary experimental feeding trials that suggested it would provide enough drift to persist across the original 48 $hr$ time period of *Sequence 1*. However, resource depletion was still a concern, particularly at relatively low levels of drift. Therefore, to enable finer-scale consumptive measurements and subsequent parameter estimation at relatively low drift levels, we initiated *Sequence 2* Trials, which halved the number of urchins and reduced the experimental duration. Our model fitting approach utilized all of these Trials, generating parameter estimates at the per-urchin scale. The urchin barrens from which our urchins were collected exhibited a density of 17.2 $m^{-2}$, and urchins had a mean gonad index of 0.645 ($std = 0.985$, $n = 49$), a very low value that reflects urchins originating from an urchin barren (Angwin *et al.*, 2022). The 4-5 $cm$ (test diameter) urchins utilized were the most common adult size class available.

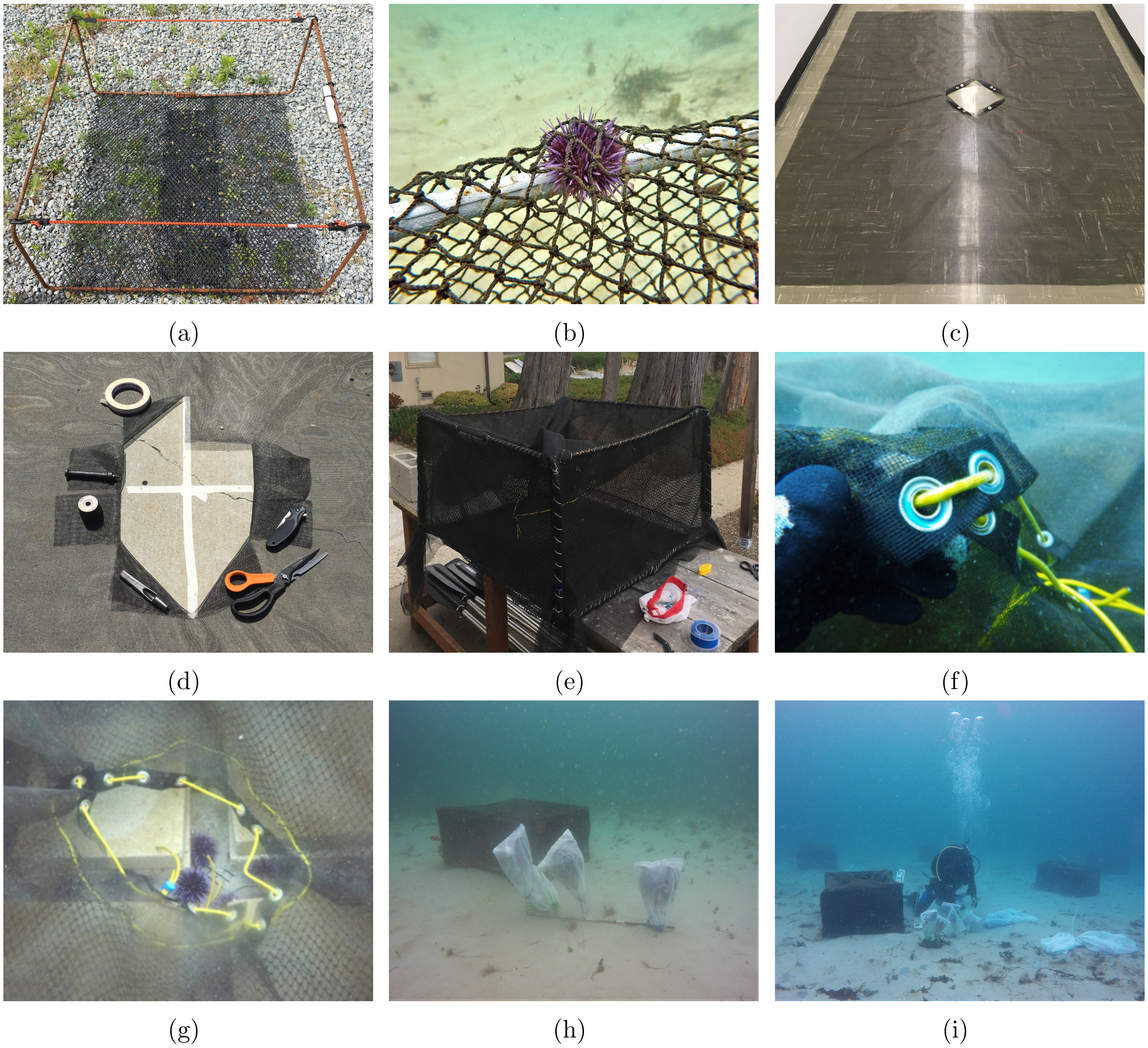

Figure S1: $(a)$ A rebar cage frame, vexar floor, flexible bungee struts, and zinc anode. $(b)$ The knotted nylon netting that was the sole layer of cage siding (summer 2018) and subsequently the interior or two layers (summer 2019). $(c)$ A sheet of fiberglass window screen and an early prototype cage "door" developed for summer 2019. $(d)$ Prepping for grommet installation to create a door through the fiberglass window screening. $(e)$ Both the nylon netting and the screening were stitched to the rebar frame. $(f)$ The grommet door frame with paracord that cinched both the netting and screening doors closed. $(g)$ The double-door access point through the top of the cage. $(h)$ 1 $m$ lengths of rebar with labeled attachments were used to organize treatments for deployment and retrieval. $(i)$ Cages were deployed in a rectangular array comprised of four rows (five cages per row), with 5 $m$ spacing between cages and rows.

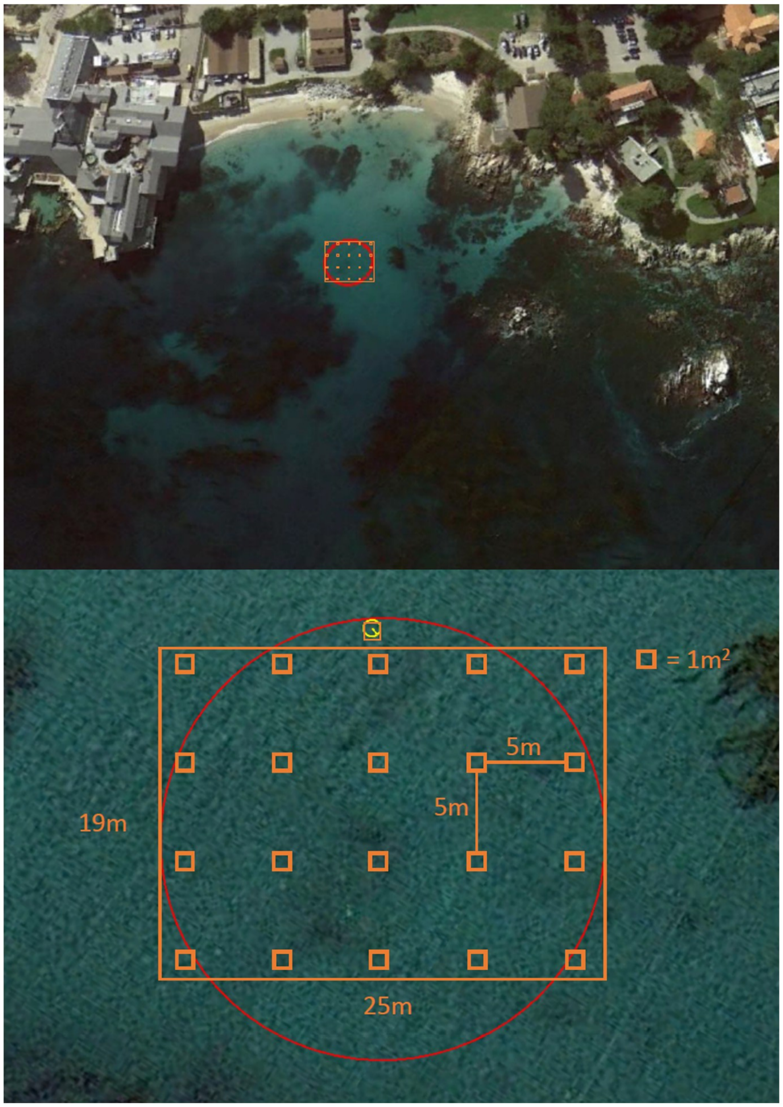

Figure S2: A representation via Google Earth of the experimental cage array located just offshore of Hopkins Marine Station, California. The array was distributed across a sandy stretch of seafloor at a depth of approximately 6 $m$, distanced at least 10 $m$ from any nearby rocky reef. The cage array formed a 25 x 19 $m$ grid with four rows of five cages; the cages were spaced uniformed spaced 5 $m$ apart from one another. This was an ideal experimental location due to its ease of access from shore via Hopkins Marine Station, its shelter from prevailing swell events out of the north or west, and the shallow soft-sediment seafloor where multiple extended dives could take place daily.

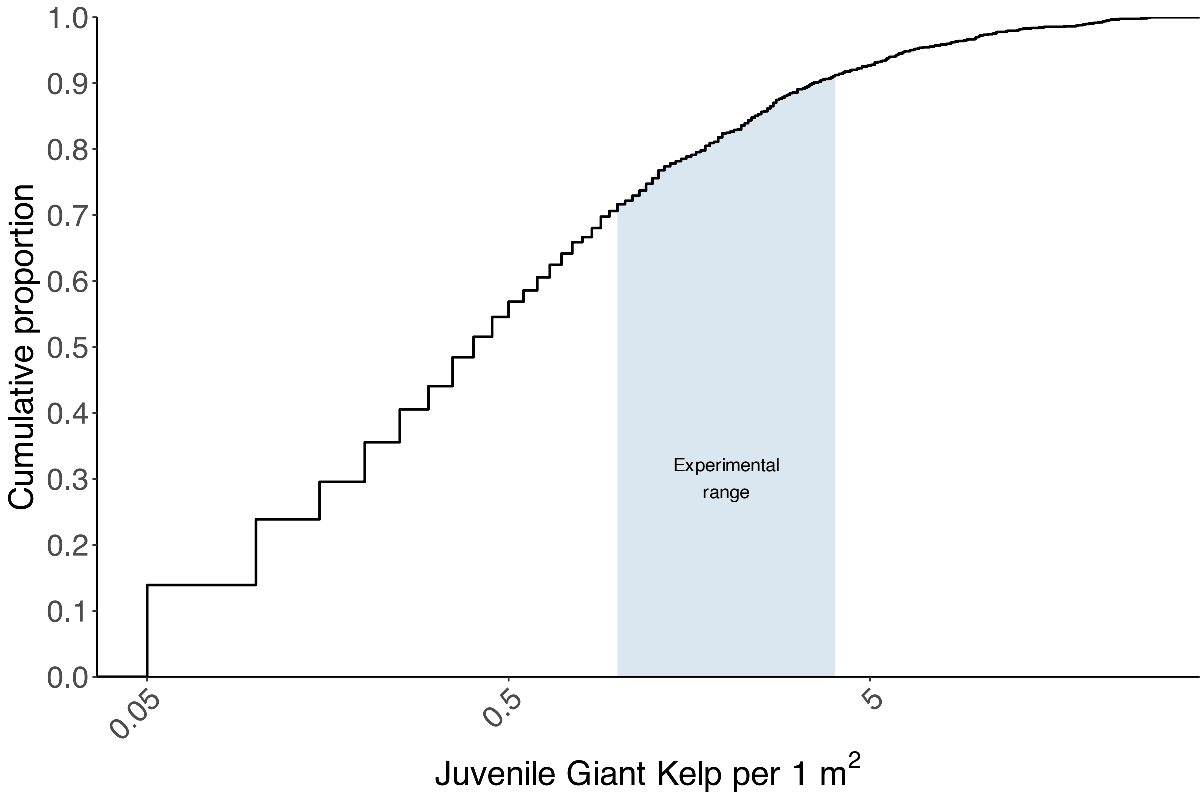

Figure S3: The cumulative frequency distribution of juvenile *Macrocystis pyrifera* kelp densities over the 38-year (1980–2018) time series of biannual monitoring around San Nicolas Island, California (Randell *et al.*, 2022), with counts of zero individuals removed. The shaded area depicts the range of densities used in our experiments (i.e., 1 and 4 individuals in each 1 $m^2$ cage). Although densities lower than our low-density treatment are common ($\sim 72\%$), densities higher than our high-density treatment are infrequent ($< 9\%$).

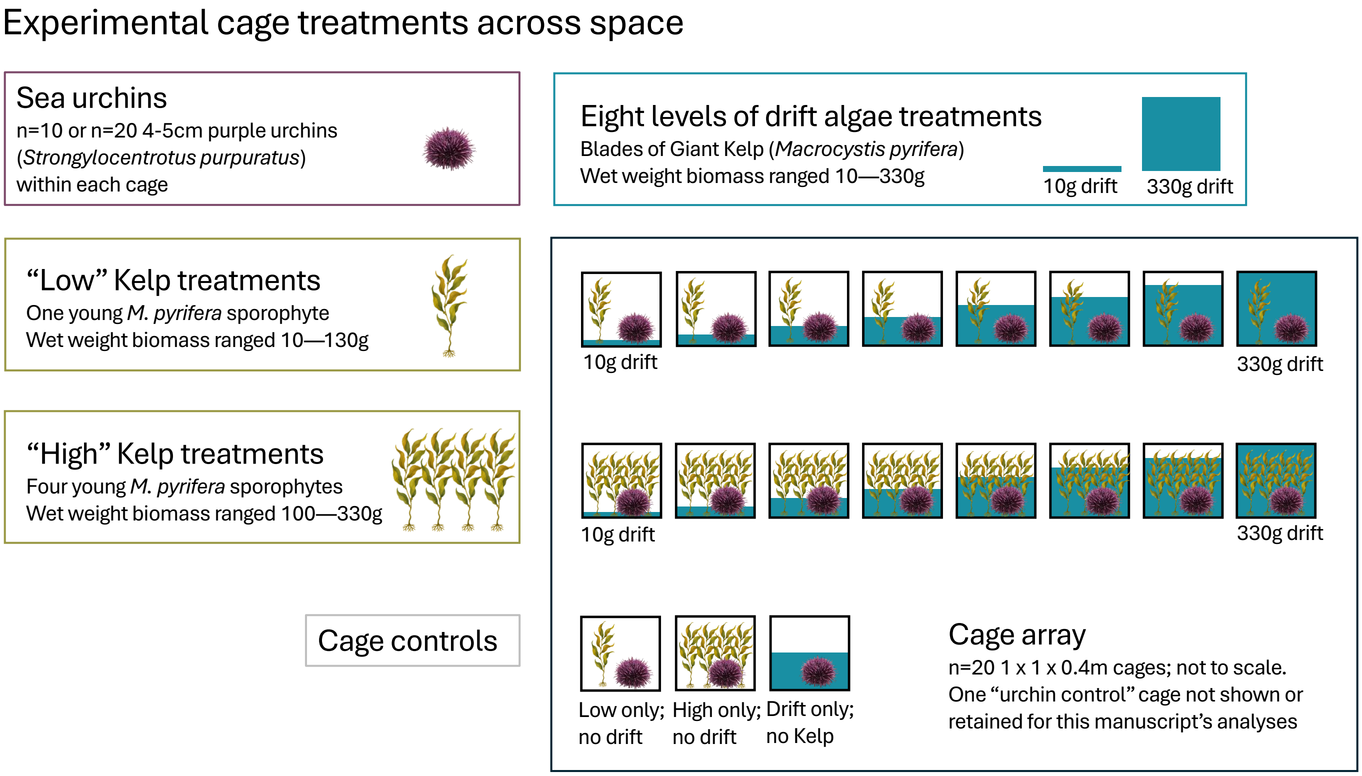

Figure S4: Our experiments consisted of 20 cages in which we imposed two live kelp treatments—Low kelp (one juvenile *M. pyrifera* sporophyte) and High kelp (four juvenile *M. pyrifera* sporophytes)—and a continuum of drift densities. They also included four cage controls.

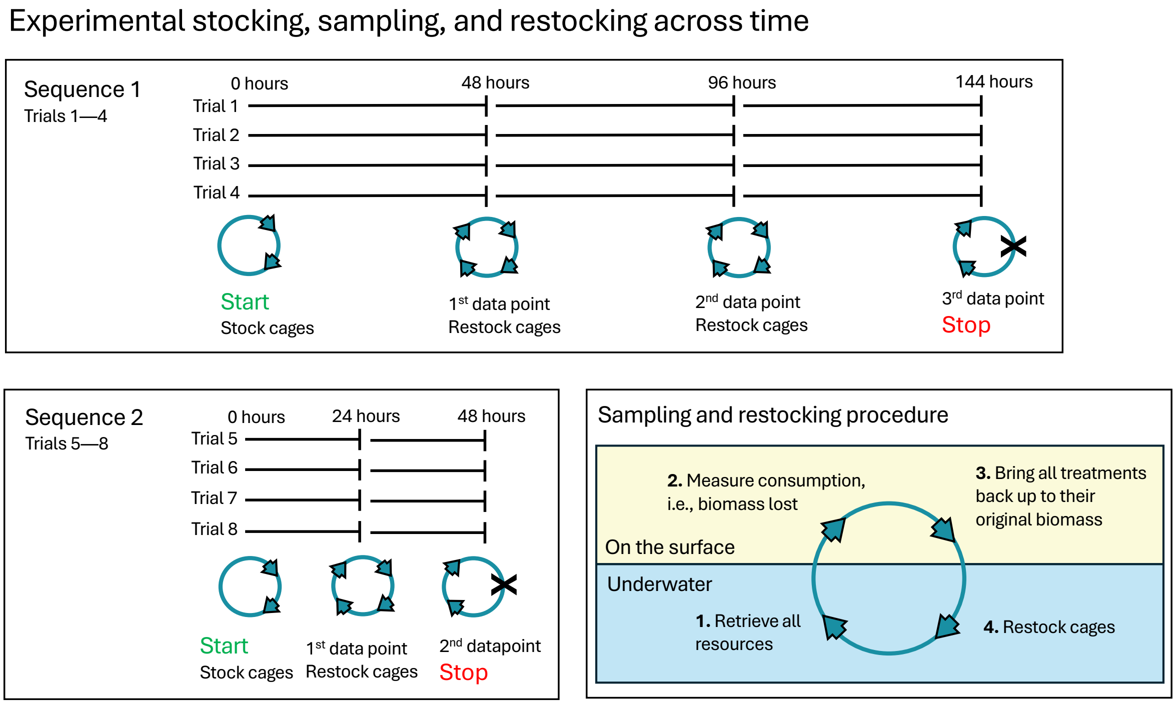

Figure S5: The temporal progression of our experiments. Each trial was an independent replicate with a new cohort of urchins and treatments randomized across cages. Resource retrieval, sampling, and restocking all took place during a single surface interval between scuba dives on a small inflatable moored above the cage array, thus ensuring a rapid turnaround for the restocking events.

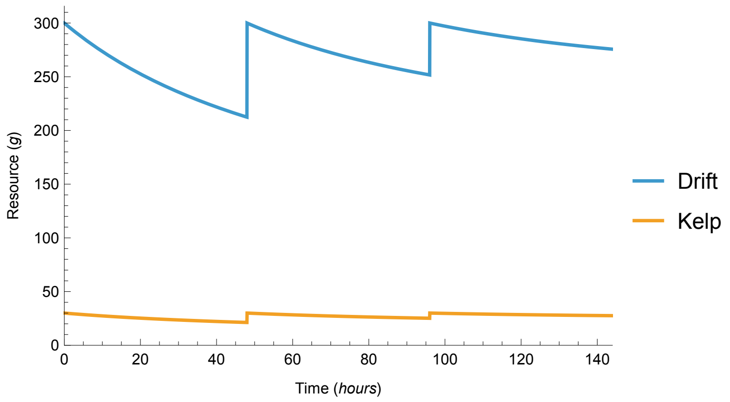

Figure S6: The idealized temporal progression of a trial, depicting the amount of drift and live kelp remaining over time. Spikes reflect our restocking of drift and kelp immediately after quantifying the amounts of each that remained. The slowing rate of decline reflects decreasing consumption by urchins.

Table S1: Sequence- and trial-specific treatment details.

| **Sequence** | **Trial** | **Time periods (*hrs*)** | **Urchins** | **Kelp** | **Replicates** |
|---|---|---|---|---|---|
| 1 | 1 | $0-44$, $44-89$, $89-134$ | 20 | 1 or 2 | 12 |
| 1 | 2 | $0-44$, $44-89$, $89-134$ | 20 | 1 or 4 | 16 |
| 1 | 3 | $0-44$, $44-89$, $89-134$ | 20 | 1 or 4 | 16 |
| 1 | 4 | $0-44$, $44-89$, $89-134$ | 20 | 1 or 4 | 15 |
| 2 | 5 | $0-16$, $16-36$ | 20 | 1 or 4 | 15 |
| 2 | 6 | $0-16$, $16-36$ | 20 | 1 or 4 | 16 |
| 2 | 7 | $0-16$, $16-36$ | 10 | 1 or 4 | 15 |
| 2 | 8 | $0-16$, $16-36$ | 10 | 1 or 4 | 16 |

#### Sequence 2 results

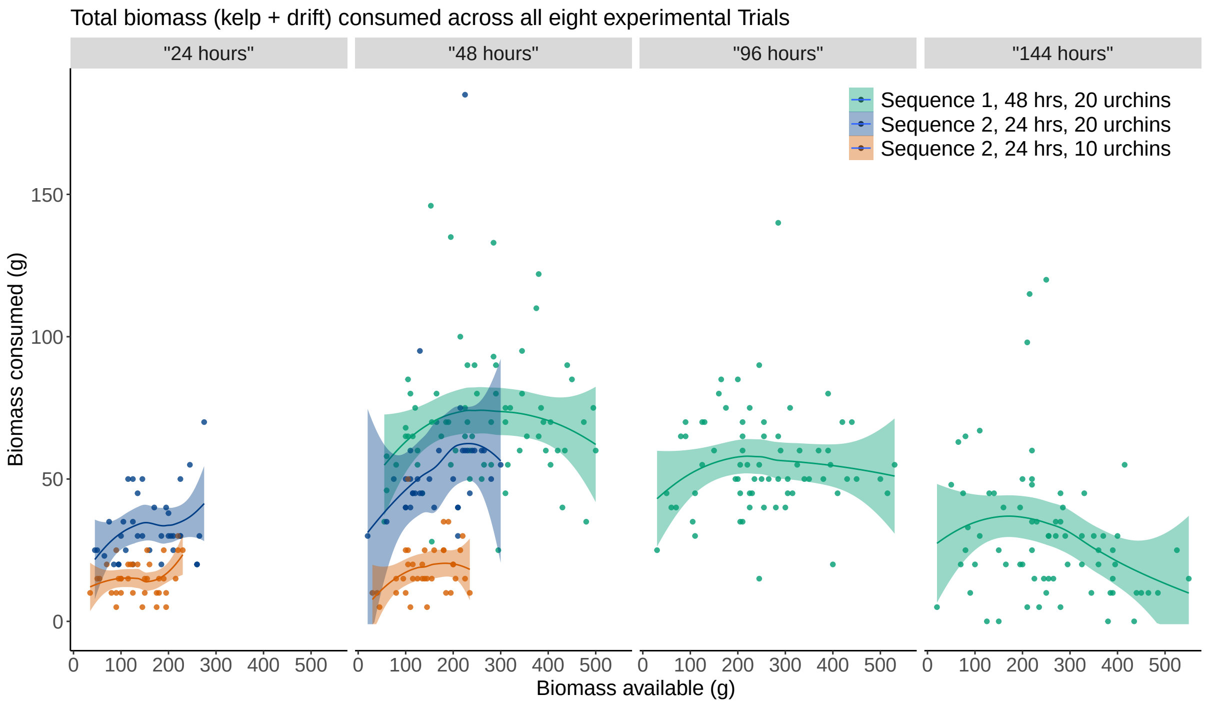

Figure S7: Total biomass of kelp and drift consumed as a function of initial biomass available. The column labels within quotations provide the approximate sample times for *Sequence 1*, the 48 hour trials (columns 2, 3, and 4) and *Sequence 2*, the 24 hour Trials (columns 1, 2).

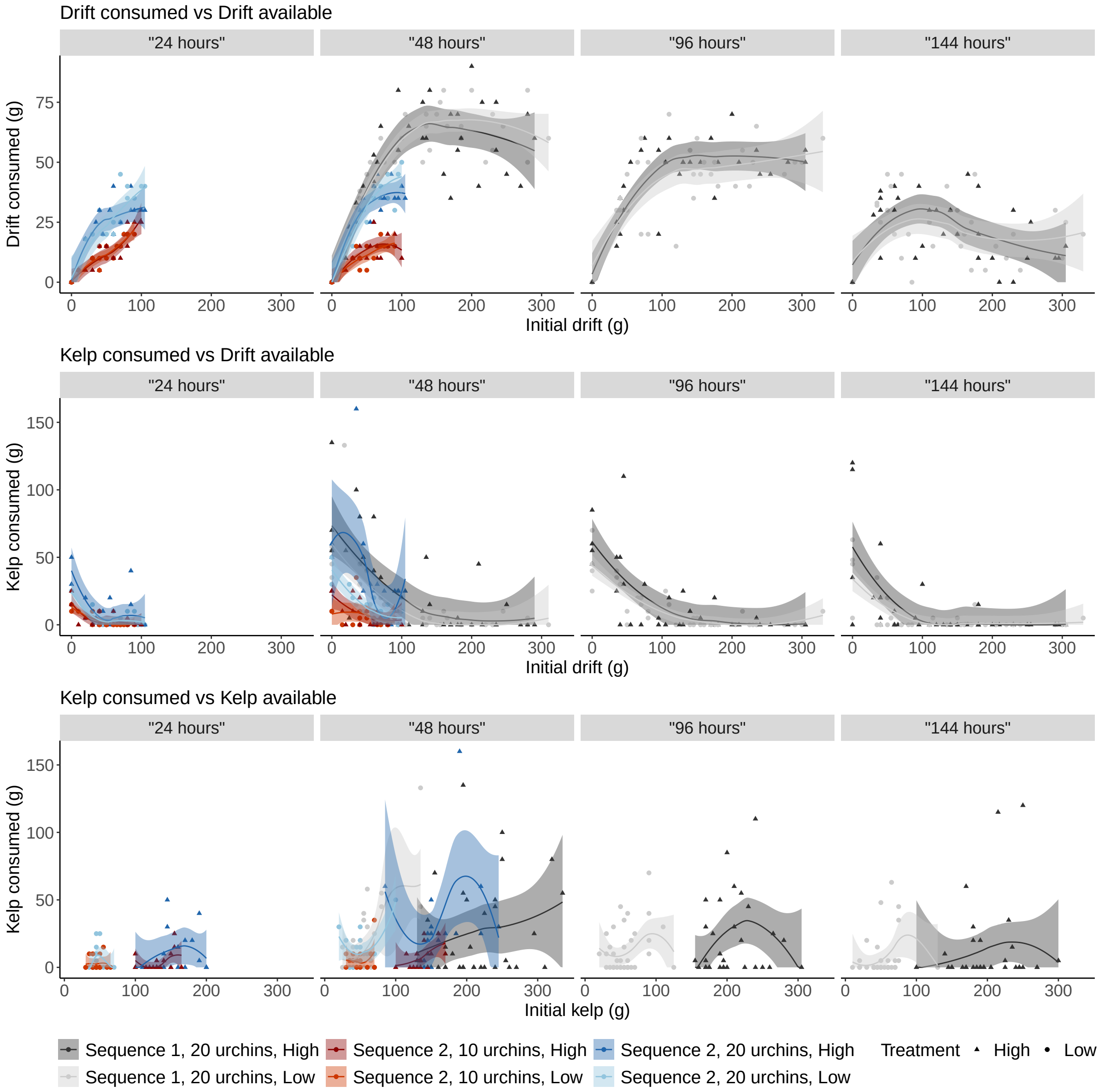

Figure S8: As in Fig. 2 of the main text, but with the data of *Sequence 2* included. The column labels within quotations provide the approximate sample times for *Sequence 1*, the 48 hour trials (columns 2, 3, and 4) and *Sequence 2*, the 24 hour Trials (columns 1, 2).

#### S2 Preference formulations

We here describe the alternative preference function formulations that we considered, doing so by demonstrating the equivalencies and differences among the published "resource-switching" functional response models in which they are embedded. These functional response models are the Yodzis model of Koen-Alonso (2007) used in the main text, a generalization of the logistic model of Elton & Greenwood (1970), and a reduced version of the model of van Leeuwen *et al.* (2013).

Focusing on just two resources ($S$ for drift, $A$ for kelp), we use subscripts to refer to a given focal resource and superscripts to refer to different models. (This notation differs from the main text where we used $p$ to refer to the urchins' preference for drift under the Yodzis model implicitly.) For example, while $p_S(S, A)$ will refer to the consumer's preference for $S$ given the availability of $S$ and $A$ in general (i.e. for no specific model or formulation), $p_S^Y(S, A)$ will refer to this preference function under the Yodzis formulation. Throughout, note that $p_S = 1 - p_A$ given how preferences are formulated.

##### Logistic and Yodzis models

We first demonstrate the mathematical equivalence of the Logistic and Yodzis formulations (because this has apparently not been previously shown in the literature). We then provide an interpretation of the parameters and explain the statistical reformulation we used in estimating the parameters.

###### Mathematical equivalence

The standard logistic function

$$f(x) = \frac{1}{1 + e^{-x}} \tag{S1}$$

due to Verhulst (1838) may be used to model how the preference of a consumer for resource $S$ increases as a function of the ratio of available $S$ and $A$ by replacing $x$ with $\log(S/A)$ such that

$$\begin{aligned} p_S(S, A) &= \frac{1}{1 + e^{-\log(S/A)}} && \text{(S2a)} \\ &= \frac{1}{1 + \left(\frac{S}{A}\right)^{-1}} && \text{(S2b)} \\ &= \frac{1}{1 + \frac{A}{S}}\,. && \text{(S2c)} \end{aligned}$$

However, because $f(x)$ has the symmetry property that $1 - f(x) = f(-x)$ and because it is easier to think about preference for $S$ as increasing with $S/A$ (rather than $A/S$), we write

$$p_S(S, A) = 1 - \frac{1}{1 + \frac{S}{A}}\,. \tag{S3}$$

When written to include an exponent parameter $\phi$ as

$$p_S(S, A) = 1 - \frac{1}{1 + \left(\frac{S}{A}\right)^{\phi}} \tag{S4}$$

we obtain the model of Elton & Greenwood (1970) and Greenwood & Elton (1979). By including a second parameter $w$, we obtain what we refer to as the logistic preference model ($L$), written as

$$p_S^L(S,A) = 1 - \frac{1}{1 + \left(\frac{w}{1-w}\right)\left(\frac{S}{A}\right)^{\phi}}\,. \tag{S5}$$

This logistic preference model is mathematically equivalent to a two-species version of the preference component of the functional response model that Koen-Alonso (2007) attributes to Peter Yodzis ($Y$):

$$p_S^Y(S,A) = \frac{wS^{\phi}}{wS^{\phi} + (1-w)A^{\phi}}\,. \tag{S6}$$

Their equivalence is shown as follows (dropping the model-specific superscript to $p_S$ in between):

$$p_S^L(S,A) = 1 - \frac{1}{1 + \left(\frac{w}{1-w}\right)\left(\frac{S}{A}\right)^{\phi}} \tag{S7a}$$

$$p_S + \frac{1}{1 + \left(\frac{w}{1-w}\right)\left(\frac{S^{\phi}}{A^{\phi}}\right)} = 1 \tag{S7b}$$

$$p_S\left(1 + \left(\frac{w}{1-w}\right)\left(\frac{S^{\phi}}{A^{\phi}}\right)\right) + 1 = 1 + \left(\frac{w}{1-w}\right)\left(\frac{S^{\phi}}{A^{\phi}}\right) \tag{S7c}$$

$$p_S\left(1 + \left(\frac{w}{1-w}\right)\left(\frac{S^{\phi}}{A^{\phi}}\right)\right) = \frac{w}{1-w}\frac{S^{\phi}}{A^{\phi}} \tag{S7d}$$

$$p_S\left(1 + \frac{w}{1-w}\frac{S^{\phi}}{A^{\phi}}\right)(1-w)A^{\phi} = wS^{\phi} \tag{S7e}$$

$$p_S\left((1-w)A^{\phi} + wS^{\phi}\right) = wS^{\phi} \tag{S7f}$$

$$p_S = \frac{wS^{\phi}}{wS^{\phi} + (1-w)A^{\phi}} = p_S^Y(S,A)\,. \tag{S7g}$$

The consumer's preference for resource $A$ differs from the above only in the numerator of the Yodzis model. Hence, due to symmetry,

$$p_A^Y(S,A) = \frac{(1-w)A^{\phi}}{wS^{\phi} + (1-w)A^{\phi}} = \frac{1}{1 + \left(\frac{w}{1-w}\right)\left(\frac{S}{A}\right)^{\phi}} = p_A^L(S,A)\,. \tag{S8}$$

Note that, despite their equivalence for two resources, the Yodzis model is easily generalized to an arbitrary number of resources (see Koen-Alonso, 2007) while the logistic model is not. Their equivalence in the two-resource case demonstrates why the consumer's preference under the Yodzis model is also function of the ratio of the two resources' abundances rather than their absolute abundances.

##### Parameter interpretations

Parameter $w$ may be interpreted as the consumer's baseline preference for $S$ relative to $A$ (i.e., its preference for resource $S$ when the two resource are equally abundant). The value of $w$ is constrained by $0 \leq w \leq 1$. It is interpreted as a proportion because in the many resources case $\sum_{i=1}^{S} w_i = 1$. The ratio $w/(1-w)$ reflects the odds of the consumer choosing $S$ over $A$ given equal resource abundances. The model of Elton & Greenwood (1970) assumes no preference at equal resource abundances (i.e., that $w = 0.5$).

Parameter $\phi$ reflects the sensitivity of the consumer's preference to changes in the relative abundance of the two resources; it is a switching sensitivity or rate (although not a rate in time). Values of $\phi > 0$ reflect positive switching (i.e., an increasing preference for $S$ as its abundance relative to the abundance of $A$ increases), values of $\phi < 0$ reflect negative switching (i.e., a decreasing preference for $S$ as its relative abundance increases), and a value of $\phi = 0$ reflects a constant density-independent preference (i.e., the preference for $S$ remains at $w$ regardless of its relative abundance).

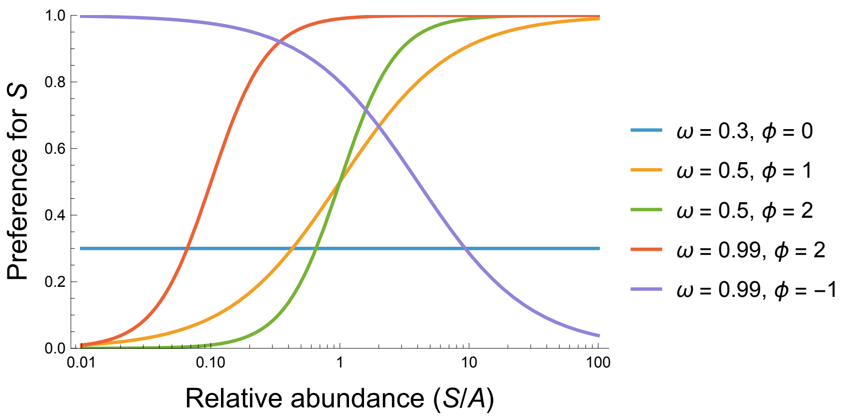

Figure S9: The generalized logistic and Yodzis models for different values of the baseline preference $w$ and the switching sensitivity $\phi$.

##### Alternative statistical formulation

A model's mathematically equivalent formulations need not be statistically equivalent. More specifically, it is possible for the statistical properties of a model's parameters to be more easily estimated under some formulations than for other mathematically equivalent formulations (e.g., the Holling and Michaelis-Menten Type II functional response formulations, see Novak & Stouffer, 2021). One way that this manifests in the context of Bayesian analyses is in the specification of parameter priors when the desire is for these to be uninformative or "weakly" informative (e.g., when there is little previous knowledge with which to constrain a prior's parameter values and insufficient signal in the data to negate the influence of a given assumed prior distribution). In such situations it is often the case that some prior distributions, such as the Gaussian distribution, are preferred over other, more constrained distributions.

Pertinent to the Yodzis and Logistic preference models, naively choosing to specify a uniform (or beta) prior on the baseline preference parameter given its interpretation as a proportion subject to $0 \leq w \leq 1$ would (or could) result in substantial probability being given to very low and very high parameter values due to the structural positioning of $w$ as an odds ratio in the model. This issue is well-recognized and discussed in the statistical literature on logistic regression. We therefore chose to fit an alternative formulation, expressing

$$p_S(S, A) = 1 - \frac{1}{1 + \exp\left[\varpi + \phi \log\left(\frac{S}{A}\right)\right]} \,. \tag{S9}$$

Although mathematically equivalent to $p_S^L$ and $p_S^Y$, this formulation permits one to place weakly informative Gaussian distributions on both $\varpi$ and $\phi$ (i.e., entirely uninformative regarding the relative preference for drift versus kelp as determined by the distribution's mean, but weakly informative regarding the degree of possible preference as determined by the distribution's standard deviation).

While the interpretation of $\phi$ remains unchanged, $\varpi$ is related to $w$ in reflecting baseline preference in terms of the log-odds (i.e., the logit link function),

$$\varpi = \log \tfrac{w}{1-w} \,. \tag{S10}$$

While proportions are constrained ($0 \leq w \leq 1$), the log-odds are unconstrained ($-\infty \leq \varpi \leq \infty$). We may thus place a Gaussian prior on $\varpi$ that is centered on an expected value of $\mu = 0$ (corresponding to $w = 0.5$ such that there is no baseline preference for either $S$ or $A$) and has a standard deviation $\sigma$ whose value is chosen to affect a near-uniform prior on $w$. Several other priors are also possible and have additional benefits in various statistical contexts. In our application, use of these alternative priors did not alter our conclusions.

##### Preference and switching point

Based on the statistical reformulation of our preference model (eqn. S9) with parameters $\varpi$ and $\phi$, the consumer's preference for $S$ when the two resources are equally abundant is

$$1 - \frac{1}{1 + \exp[\varpi]} = \frac{1}{1 + \exp[-\varpi]} \,. \tag{S11}$$

The "switching point" ratio of resource abundances where the consumer's preferences for the two resources are equal is

$$\exp\left[\frac{\varpi}{\phi}\right] \,. \tag{S12}$$

#### Reduced van Leeuwen et al. model

van Leeuwen *et al.* (2013) derived a functional response model for a consumer that switches between an arbitrary number of resource with preferences that depend on the resource it consumed last. Their model may be written as

$$f_i = \frac{a_i N_i \sum_{k=1}^{n} s_{ik} a_k N_k}{\sum_{k=1}^{n} a_k N_k \left(1 + \sum_{j=1}^{n} s_{kj} h_{kj} a_j N_j\right)} \,, \tag{S13}$$

and reduces to multi-species Holling Type II response (with resources abundances $N$, attack rates $a$, and handling times $h$) when all "switching similarities" $s$ equal 1 (whereby realized attack rates and handling times are independent of the previously consumed resource). By focusing on the preference component, their model can be reduced to a two-species preference model analogous to $p^L$ and $p^Y$ discussed above. Specifically, by letting $h_i = 0$, $s_{ii} = 1$, and $s_{ij} = s_{ji} = \psi^{-1}$, substituting $\nu = a_S/a_A$, and expressing the preference for drift as $p_S = \frac{f_S}{f_S+f_A}$, one can obtain

$$p_S^V(S,A) = 1 - \frac{1+\frac{\nu}{\psi}\frac{S}{A}}{1+2\frac{\nu}{\psi}\frac{S}{A}+\left(\nu\frac{S}{A}\right)^2} = \frac{\nu\left(1+\nu\psi\frac{S}{A}\right)\frac{S}{A}}{\psi+\nu\left(2+\nu\psi\frac{S}{A}\right)\frac{S}{A}} \tag{S14}$$

wherein $\nu$ reflects the baseline preference for $S$ over $A$ (like the the odds ratio $\exp(\varpi)$) and $\psi$ reflects a switching rate similar to $\phi$ above (Fig. S10).

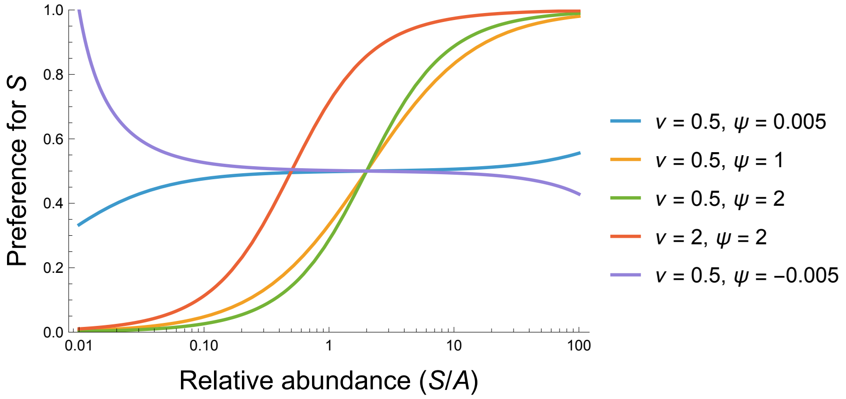

Figure S10: The reduced van Leeuwen model for different values of the baseline preference parameter $\nu$ and the switching rate parameter $\psi$.

##### Alternative statistical formulation

Motivated by the same reasoning as for the logistic model, we reformulate the reduced van Leeuwen model by substituting $\beta = \log(\nu/\psi)$ and $\gamma = \log(\nu^2)$ and expressing the resources abundances as a log ratio to obtain

$$p_S^V(S,A) = 1 - \frac{1+\exp\left[\beta+\log\left(\frac{S}{A}\right)\right]}{1+\exp\left[\log(2)+\beta+\log\left(\frac{S}{A}\right)\right]+\exp\left[\gamma+2\log\left(\frac{S}{A}\right)\right]}. \tag{S15}$$

After fitting this model to the data with weakly-informative zero-centered Gaussian priors on $\beta$ and $\gamma$, we then back-solve for the baseline preference $\nu$ and the switching rate $\psi$.

##### Preference and switching point

Based on the statistical reformulation of the reduced VanLeeuwen preference model (eqn. S15) with parameters $\beta$ and $\gamma$, the consumer's preference for $S$ when the two resources are equally abundant is

$$1 - \frac{1 + \exp[\beta]}{1 + \exp[\log(2) + \beta] + \exp[\gamma]} = \frac{\exp[\beta] + \exp[\gamma]}{1 + 2\exp[\beta] + \exp[\gamma]} \,, \tag{S16}$$

while the "switching point" ratio of resource abundances where the consumer's preferences for the two resources are equal is simply

$$\exp\left[0.5\gamma\right] \,. \tag{S17}$$

### S3 Movement suppression

We here provide two independent mechanistic derivations of the movement suppression function used in the main text,

$$m := m(S, A) = \frac{1}{1 + ab\,E^2} \,, \tag{S18}$$

where $E := pS + (1 - p)A$ denotes the preference-weighted effective resource density and $b$ is the movement suppression parameter. While equation S18 is structurally similar to the Monod-Haldane type IV functional response, it does not describe substrate inhibition or resource toxicity. Rather, as we show below, it arises from assumptions regarding the density-dependent behavior of urchins that was directly observable in our experiments.

#### Time-budget derivation

We first derive the movement function by expressing the urchin's time as a budget between active searching and stationary foraging.

##### Encounter rate when active and passive

While actively moving, each urchin encounters the effective resource density at rate

$$\lambda = aE \,, \tag{S19}$$

where $a$ is the search rate. Upon encountering resource the urchin stops to consume, incurring a per-item consumption ("handling") time $\tau_0$. In the standard type II functional response, the consumer resumes searching immediately after each "handled" item (Novak *et al.*, 2025). For urchins in our experiment, however, an additional process was apparent: while stationary and consuming, they continued to passively encounter nearby resources. We assume this occurs at the same rate $\lambda$, with each additional encounter extending the stationary time by a further expected time $\tau_0$ before the urchin considers returning to active movement. We assume urchins return to active movement at a background rate $\gamma$ that is independent of resource density. The expected number of passive re-encounters before resuming movement is therefore $\lambda/\gamma = aE/\gamma$, yielding a total expected time per initial encounter of

$$\tau_{\text{eff}}(E) = \tau_0 \left(1 + \frac{aE}{\gamma}\right) \,. \tag{S20}$$

Note that $\tau_{\text{eff}}$ is a linear function of $E$: the expected time an urchin remains passive increases with resource density because denser resources sustain passive re-encounters before the urchin would otherwise resume active movement.

##### Movement fraction

Let $T_s$ denote total time spent actively searching during a period of duration $T$. The expected number of initial encounters is $N_e = aE \cdot T_s$, so total time is $T = T_s + N_e \cdot \tau_{\text{eff}}$, or

$$T = T_s \left( 1 + a\tau_0 E + \frac{a^2 \tau_0}{\gamma} E^2 \right), \tag{S21}$$

giving a fraction of time spent actively moving of

$$m = \frac{T_s}{T} = \frac{1}{1 + a\tau_0 E + \dfrac{a^2 \tau_0}{\gamma} E^2}. \tag{S22}$$

The quadratic term in the denominator of eqn. S22 arises from the compounding of two linear dependencies on resource density: the initial encounter rate while searching and the expected additional time spent stationary per encounter. This structure is formally analogous to the Monod-Haldane expression, but with the density-dependent term arising from stationary behavior rather than substrate inhibition.

In the regime where passive re-encounters while stationary are frequent relative to the return-to-movement rate such that $aE/\gamma \gg 1$, the quadratic term in the denominator dominates the linear term such that

$$m \approx \frac{1}{1 + \dfrac{a^2 \tau_0}{\gamma} E^2}. \tag{S23}$$

Defining $b \equiv a\tau_0/\gamma$ then recovers eqn. S18. Parameter $b$ may therefore be interpreted as the ratio of search rate $\times$ per-item handling time to the resumption-of-movement rate, with large values of $b$ reflecting urchins that rarely leave the stationary state.

#### Behavioral-state derivation

The same form may be derived by treating urchin behavioral modes as a continuous-time Markov chain.

##### States and transitions

We define three mutually exclusive behavioral states. State M (actively moving) corresponds to urchins moving at full velocity and encountering resources. State T (tentatively settled) corresponds to urchins that have made first contact with resources and slowed, but that may still resume movement without having committed to stationary behavior. State C (committed foraging) corresponds to urchins that have fully settled into a crevice such that movement is suppressed. The distinction between T and C is consistent with our direct observations: at modest drift densities urchins contacted drift but often resumed movement, whereas at higher drift densities urchins wedged themselves into the paving-stone crevices and remained there for the duration of the experiments.

Both forward transitions require an encounter event and therefore scale with $E$,

$$\text{M} \xrightarrow{aE} \text{T} \xrightarrow{aE} \text{C}, \tag{S24}$$

while the reverse transitions occur at rates $\gamma_1$ (T → M) and $\gamma_2$ (C → T), with $\gamma_1 \gg \gamma_2$. That is, tentatively settled urchins often resume movement, whereas committed feeders rarely do.

##### Steady-state solution

Denoting the fraction of time spent in each state as $\pi_M$, $\pi_T$, and $\pi_C$, respectively, and setting $d\pi_T/dt = 0$ and $d\pi_C/dt = 0$, the frequencies may be expressed in terms of $\pi_M$ as

$$\pi_T = \frac{aE}{\gamma_1 + aE}\,\pi_M \tag{S25a}$$

$$\pi_C = \frac{aE}{\gamma_2}\,\pi_T = \frac{a^2E^2}{\gamma_2(\gamma_1 + aE)}\,\pi_M. \tag{S25b}$$

Since $\pi_M + \pi_T + \pi_C = 1$, the fraction of time spent in active movement will be

$$m = \pi_M = \frac{\gamma_2(\gamma_1 + aE)}{\gamma_1\gamma_2 + 2\gamma_2 aE + a^2E^2}\,. \tag{S26}$$

In the limit $\gamma_1 \gg aE$ (i.e., the tentative phase is brief; urchins typically resume active movement after a first encounter), eqns. S25a–S25b reduce to $\pi_T \approx (aE/\gamma_1)\,\pi_M$ and $\pi_C \approx (a^2E^2/\gamma_1\gamma_2)\,\pi_M$, and eqn. S26 reduces to

$$m \;\approx\; \frac{1}{1 + \dfrac{a^2}{\gamma_1\gamma_2}\,E^2}\,. \tag{S27}$$

Defining $b \equiv a/(\gamma_1\gamma_2)$ recovers eqn. S18 assumed in the main text. Parameter $b$ may be interpreted as the search rate divided by the product of the two behavioral departure rates, with large values of $b$ reflecting rare transitions back to active movement from either settled state.

##### Why an exponent of 2?

Both derivations show that the exponent of 2 in eqn. S18 arises from the assumption that committed settlement be preceded by two sequential density-dependent events: a first encounter while moving (rate $\propto E$) and a second encounter (sufficient passive re-exposure) before the urchin returns to active movement (probability $\propto E$). The probability of completing this two-step process therefore scales as $E^2$. More generally, if committed settlement required $n$ sequential encounters, the exponent would equal $n$.

#### S4 Model fitting, comparison, and inferences

##### Bayesian model fitting

We used a Bayesian approach to fit eight competing models to the data using *Stan* v.2.32.7. (Stan Development Team, 2019) and *cmdstanr* v.0.9.0 in *R* v.4.5.1. The models all had the basic structure of eqns. 1 and 2 of the main text, but differed from each other in assuming either (*i*) the logistic or the reduced van Leeuwen preference function (see *Section S2*), (*ii*) movement suppression at high resource abundances (i.e. $b > 0$ as estimated free parameter) or the absence thereof (i.e. $b = 0$ such that $m = 1$ regardless of resource abundances), and (*iii*) a gut evacuation rate ($z$ as a free parameter) or the absence thereof ($z = 0$). (We included the last of these anticipating insufficient information in the data to estimate a gut evacuation rate, the linear approximation to which could be effectively subsumed by our modeling of gut fullness sensitivity via $v$ of the hunger level $h$ process.)

The initial conditions of kelp $A[0]$ and drift $S[0]$ were the supplied experimental values for the first treatment and subsequent restocking events. The initial condition of gut fullness $F[0]$ was set to 0 because there was insufficient information content in our data to estimate it as a free parameter. The value of gut fullness at the end of period 1 was used as the initial condition of period 2. Likewise, the gut fullness at the end of period 2 was used as the initial condition of period 3. Our fitting thereby accounted for the restocking of kelp and drift at the beginning of each time period, their consumption within each time period, and allowed gut fullness to increase in a cumulative fashion across the time periods. Integration of each system of ODEs was performed using the Runga-Kutta *ode_rk45* method.

Drift and kelp weights were represented with a Gaussian likelihood. Parameter prior distributions were specified as given in Table S3 (see *Section S2* for justifications and interpretations). Four Markov chains were run for $5,000$ iterations (including a warm-up of $1,000$ iterations) with a target acceptance statistic of 0.8. The chains were well-mixed and no divergent transitions or other sampling issues were incurred except in a few warm-up iterations of some models that permitted no movement reduction ($m = 1$).

#### Model comparisons and inferences

We used cross-validation to assess the relative performance of our competing models, comparing models on the basis of their leave-one-out estimates of the expected log pointwise predictive density (Sivula *et al.*, 2025; Vehtari *et al.*, 2025). Models weights were determined using the pseudo-BMA+ method (Yao *et al.*, 2018).

Table S2: Relative model performance as assessed by the Bayesian LOO estimate of the expected log pointwise predictive density. $p$ is the effective number of parameters. Model weights estimated using the pseudo-BMA+ method. Models ordered by weight. A parenthetical $z$ indicates a model including a gut evacuation rate, assumed absent in other models. A parenthetical $m = 1$ indicates a model with $m$ fixed to 1 (equivalent to $b = 0$), meaning no movement suppression at high resource abundances.

| Model | ELPD | $SE_{ELPD}$ | $p$ | ΔELPD | $SE$(ΔELPD) | Weight |
|---|---|---|---|---|---|---|
| van Leeuwen | -2966 | 61 | 18.3 | 0.0 | 0.0 | 0.39 |
| Logistic | -2968 | 62 | 20.9 | -1.2 | 2.4 | 0.23 |
| van Leeuwen ($z$) | -2967 | 61 | 18.7 | -0.6 | 0.2 | 0.22 |
| Logistic ($z$) | -2968 | 62 | 21.2 | -1.5 | 2.5 | 0.16 |
| van Leeuwen ($m = 1$) | -3096 | 48 | 15.1 | -130.0 | 27.0 | 0.00 |
| Logistic ($m = 1$) | -3099 | 49 | 19.2 | -132.7 | 27.6 | 0.00 |
| Logistic ($z$, $m = 1$) | -3056 | 52 | 16.5 | -89.7 | 19.4 | 0.00 |
| van Leeuwen ($z$, $m = 1$) | -3051 | 52 | 14.2 | -85.1 | 18.1 | 0.00 |

Table S3: Parameter bounds, prior specifications, and posterior median estimates with 95% credible intervals. See Table S2 for model name interpretations.

| Model | Parameter | Prior | Bounds | Posterior median | CI |
|---|---|---|---|---|---|
| Logistic | Search rate ($a$) | Exp(10) | [0, 0.05] | 0.0040 | (0.0033—0.0050) |
| | Supression rate ($b$) | Exp(10) | [0, 0.1] | 0.0476 | (0.0410—0.0548) |
| | Baseline preference ($\tilde{\omega}$) | $\mathcal{N}(0, 1.8)$ | [-6, 6] | 2.7500 | (2.1720—3.6160) |
| | Switching sensitivity ($\varphi$) | $\mathcal{N}(0, 10)$ | [-5, 5] | 0.6567 | (0.2629—1.2180) |
| | Stomach sensitivity ($v$) | Exp(1) | [0, 0.5] | 0.1505 | (0.1301—0.1736) |
| | Variance ($\sigma$) | Exp(0.1) | [10, 20] | 14.7900 | (14.0300—15.6200) |
| Logistic ($m = 1$) | Search rate ($a$) | Exp(10) | [0, 0.05] | 0.0016 | (0.0013—0.0019) |
| | Baseline preference ($\tilde{\omega}$) | $\mathcal{N}(0, 1.8)$ | [-6, 6] | 2.7000 | (1.7220—4.3690) |
| | Switching sensitivity ($\varphi$) | $\mathcal{N}(0, 10)$ | [-5, 5] | 0.9938 | (0.2123—2.2100) |
| | Stomach sensitivity ($v$) | Exp(1) | [0, 0.5] | 0.2950 | (0.2526—0.3446) |
| | Variance ($\sigma$) | Exp(0.1) | [10, 20] | 17.7400 | (16.8400—18.7100) |
| Logistic ($z$) | Search rate ($a$) | Exp(10) | [0, 0.05] | 0.0040 | (0.0033—0.0049) |
| | Supression rate ($b$) | Exp(10) | [0, 0.1] | 0.0475 | (0.0408—0.0546) |
| | Baseline preference ($\tilde{\omega}$) | $\mathcal{N}(0, 1.8)$ | [-6, 6] | 2.7540 | (2.1740—3.6330) |
| | Switching sensitivity ($\varphi$) | $\mathcal{N}(0, 10)$ | [-5, 5] | 0.6590 | (0.2678—1.2230) |
| | Stomach sensitivity ($v$) | Exp(1) | [0, 0.5] | 0.1521 | (0.1312—0.1766) |
| | Stomach clearance ($z$) | $\mathcal{N}(0, 10)$ | [-14, 1] | -10.3200 | (-13.7800—-6.9100) |
| | Variance ($\sigma$) | Exp(0.1) | [10, 20] | 14.7900 | (14.0200—15.6300) |
| Logistic ($z$, $m = 1$) | Search rate ($a$) | Exp(10) | [0, 1] | 0.2260 | (0.0624—0.6124) |
| | Baseline preference ($\tilde{\omega}$) | $\mathcal{N}(0, 1.8)$ | [-6, 6] | 2.5340 | (1.7480—4.2600) |
| | Switching sensitivity ($\varphi$) | $\mathcal{N}(0, 10)$ | [-5, 5] | 0.5216 | (-0.0672—1.6950) |
| | Stomach sensitivity ($v$) | Exp(1) | [0, 8] | 4.1710 | (2.9900—5.4780) |
| | Stomach clearance ($z$) | $\mathcal{N}(0, 10)$ | [-8, 1] | -3.4340 | (-3.6700—-3.1850) |

*Continued on next page*

*Table S3 continued from previous page*

| Model | Parameter | Prior | Bounds | Posterior median | CI |
|---|---|---|---|---|---|
| | Variance ($\sigma$) | Exp(0.1) | [10, 20] | 16.7600 | (15.9100—17.7100) |
| van Leeuwen | Search rate ($a$) | Exp(1) | [0, 0.01] | 0.0039 | (0.0032—0.0048) |
| | Supression rate ($b$) | Exp(0.1) | [0, 0.1] | 0.0477 | (0.0411—0.0547) |
| | Baseline preference ($\tilde{\omega}$) | $\mathcal{N}(0, 10)$ | [-10, 30] | 4.8940 | (0.8342—14.3200) |
| | Switching sensitivity ($\varphi$) | $\mathcal{N}(0, 10)$ | [2, 20] | 7.9970 | (5.1130—17.3400) |
| | Stomach sensitivity ($v$) | Exp(0.1) | [0, 0.5] | 0.1524 | (0.1316—0.1751) |
| | Variance ($\sigma$) | Exp(0.1) | [10, 20] | 14.8000 | (14.0400—15.6400) |
| van Leeuwen ($m = 1$) | Search rate ($a$) | Exp(1) | [0, 0.01] | 0.0015 | (0.0013—0.0018) |
| | Baseline preference ($\tilde{\omega}$) | $\mathcal{N}(0, 10)$ | [-10, 30] | -0.6082 | (-9.4230—6.2140) |
| | Switching sensitivity ($\varphi$) | $\mathcal{N}(0, 10)$ | [2, 15] | 4.3230 | (3.7200—8.6900) |
| | Stomach sensitivity ($v$) | Exp(0.1) | [0, 0.5] | 0.2979 | (0.2558—0.3481) |
| | Variance ($\sigma$) | Exp(0.1) | [10, 20] | 17.7300 | (16.8600—18.7200) |
| van Leeuwen ($z$) | Search rate ($a$) | Exp(1) | [0, 0.05] | 0.0039 | (0.0032—0.0048) |
| | Supression rate ($b$) | Exp(0.1) | [0, 0.1] | 0.0476 | (0.0410—0.0549) |
| | Baseline preference ($\tilde{\omega}$) | $\mathcal{N}(0, 10)$ | [-10, 30] | 4.7830 | (0.7396—13.8400) |
| | Switching sensitivity ($\varphi$) | $\mathcal{N}(0, 10)$ | [2, 20] | 7.8990 | (5.0950—16.9000) |
| | Stomach sensitivity ($v$) | Exp(0.1) | [0, 0.5] | 0.1536 | (0.1322—0.1773) |
| | Stomach clearance ($z$) | $\mathcal{N}(0, 10)$ | [-16, 1] | -10.9100 | (-15.6100—-7.0640) |
| | Variance ($\sigma$) | Exp(0.1) | [10, 20] | 14.8100 | (14.0500—15.6400) |
| van Leeuwen ($z$, $m = 1$) | Search rate ($a$) | Exp(1) | [0, 5] | 1.9950 | (0.4425—4.5830) |
| | Baseline preference ($\tilde{\omega}$) | $\mathcal{N}(0, 10)$ | [-10, 30] | -0.1284 | (-9.3000—5.0560) |
| | Switching sensitivity ($\varphi$) | $\mathcal{N}(0, 10)$ | [2, 15] | 5.1360 | (4.4160—8.1430) |
| | Stomach sensitivity ($v$) | Exp(0.1) | [0, 10] | 6.5590 | (4.9400—8.3290) |

*Continued on next page*

*Table S3 continued from previous page*

| Model | Parameter | Prior | Bounds | Posterior median | CI |
|---|---|---|---|---|---|
| | Stomach clearance ($z$) | $\mathcal{N}(0, 10)$ | [-8, 1] | -3.2810 | (-3.5150—-3.0390) |
| | Variance ($\sigma$) | Exp(0.1) | [10, 20] | 16.6700 | (15.7400—17.6400) |

Table S4: Model-specific median posterior estimates (and 95% credible intervals) of the relative abundance of drift versus kelp at which urchins preference for the two resources is equal (expressed as $g$ of kelp per 1 $g$ of drift), and of their baseline preference (expressed as proportional preference and log-odds of consumption) for drift over kelp when the abundance of the two resources is equal. Models ordered and averaged by pseudo-BMA+ weight. See Table S2 for model name interpretations.

| Model | Switch point | Baseline preference | Baseline log-odds |
|---|---|---|---|
| Model average | 55.9 (14.1, 5624.4) | 0.96 (0.91, 0.98) | 3.1 (2.3, 3.9) |
| van Leeuwen | 54.5 (12.9, 5813.9) | 0.96 (0.94, 0.98) | 3.2 (2.8, 4.1) |
| Logistic | 65.7 (17.9, 4450.8) | 0.94 (0.90, 0.97) | 2.7 (2.2, 3.6) |
| van Leeuwen ($z$) | 51.9 (12.8, 4667.4) | 0.96 (0.94, 0.98) | 3.2 (2.8, 4.1) |
| Logistic ($z$) | 65.5 (17.7, 4273.0) | 0.94 (0.90, 0.97) | 2.8 (2.2, 3.6) |
| van Leeuwen ($m = 1$) | 8.7 (6.4, 77.1) | 0.98 (0.92, 0.99) | 3.7 (2.4, 4.5) |
| Logistic ($m = 1$) | 15.1 (6.6, 2411.8) | 0.94 (0.85, 0.99) | 2.7 (1.7, 4.4) |
| Logistic ($z$, $m = 1$) | 94.1 (0.0, $5.5 \times 10^{16}$) | 0.93 (0.85, 0.99) | 2.5 (1.7, 4.3) |
| van Leeuwen ($z$, $m = 1$) | 13.0 (9.1, 58.6) | 0.99 (0.95, 1.00) | 4.3 (3.0, 5.3) |

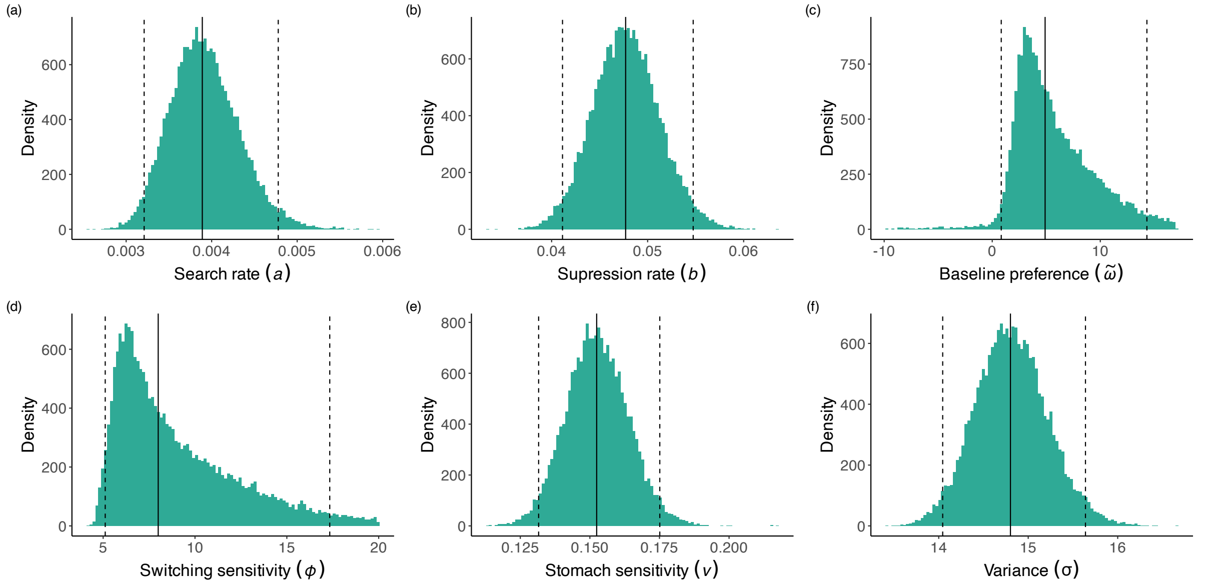

Figure S11: Posterior parameter distributions for the van Leeuwen model. The solid and dashed vertical lines respectively depict the median point estimate and 95% credible interval.

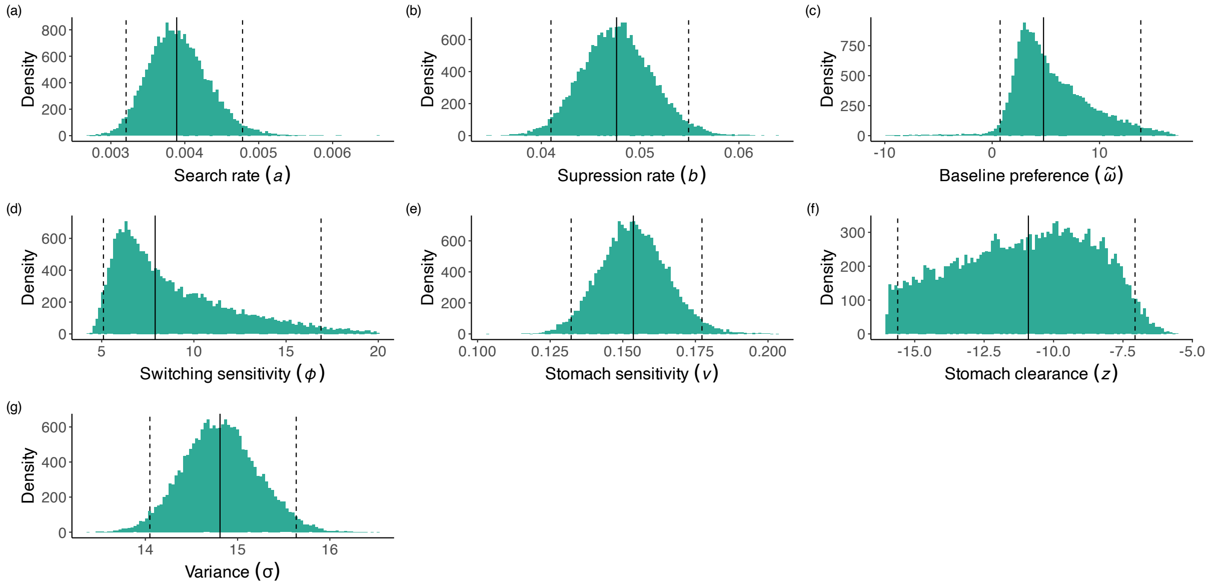

Figure S12: Posterior parameter distributions for the van Leeuwen model with gut evacuation rate ($z$). Note the logarithmic scale for $z$. The solid and dashed vertical lines respectively depict the median point estimate and 95% credible interval.

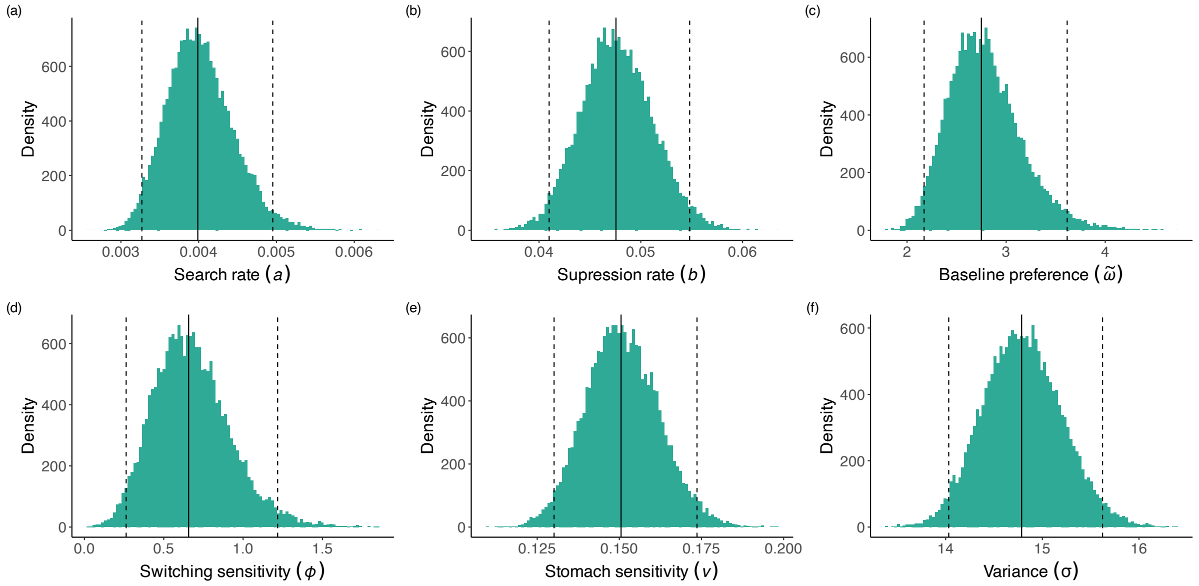

Figure S13: Posterior parameter distributions for the logistic model. The solid and dashed vertical lines respectively depict the median point estimate and 95% credible interval.

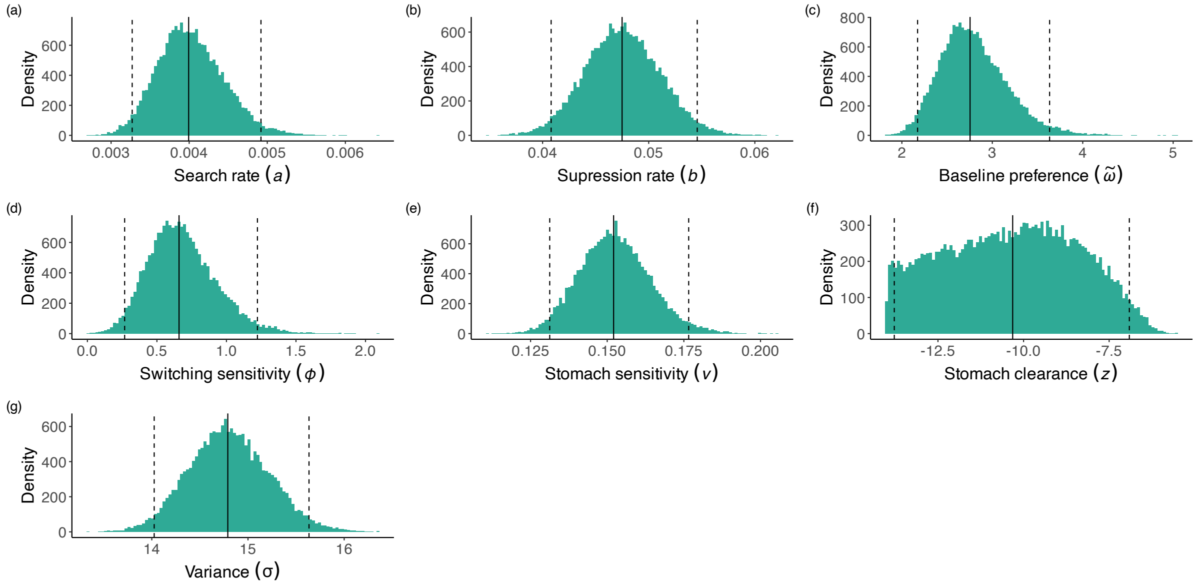

Figure S14: Posterior parameter distributions for the logistic model with gut evacuation rate ($z$). Note the logarithmic scale for $z$. The solid and dashed vertical lines respectively depict the median point estimate and 95% credible interval.

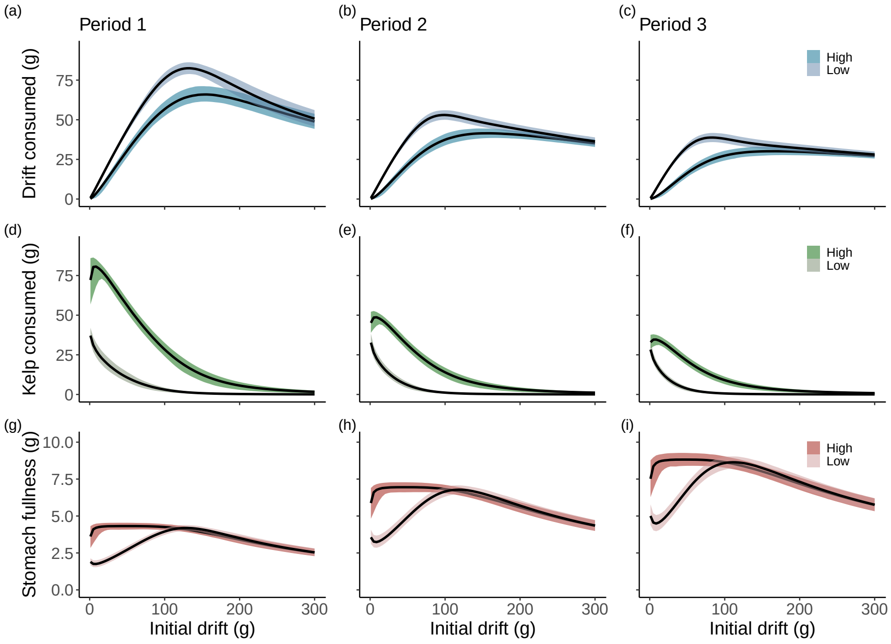

Figure S15: As in Fig. 3 of the main text for the reduced van Leeuwen preference function.

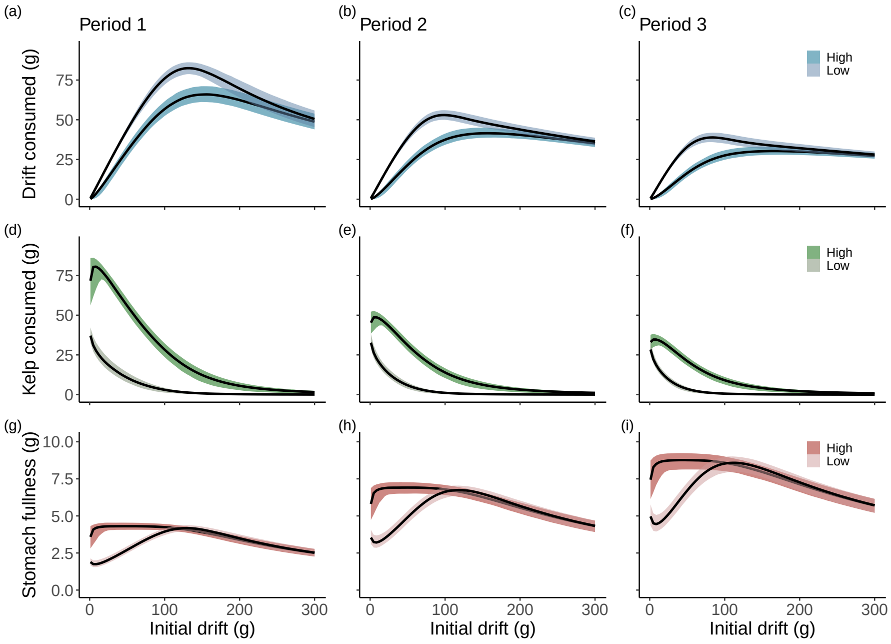

Figure S16: As in Fig. 3 of the main text for the reduced van Leeuwen preference function with gut clearance ($z$).

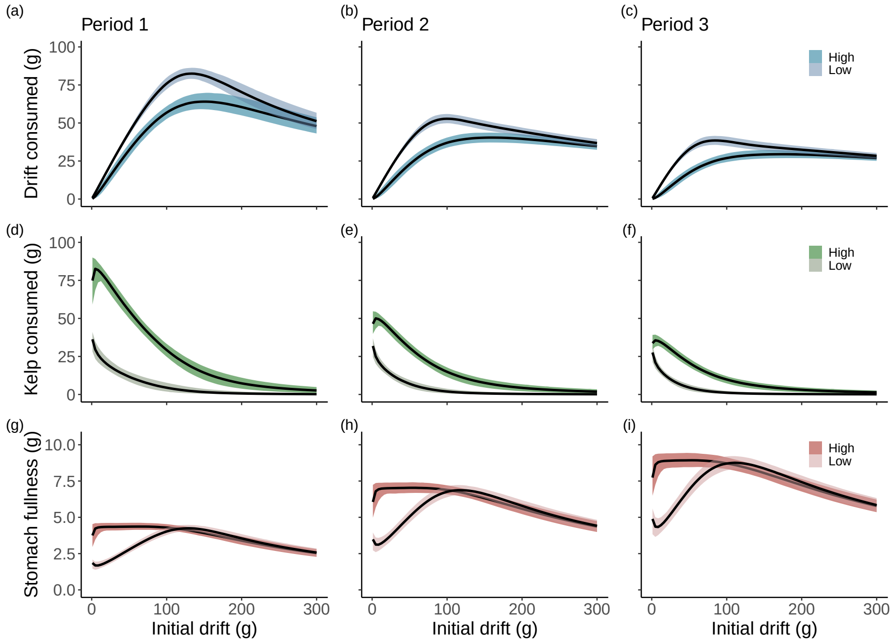

Figure S17: As in Fig. 3 of the main text for the logistic preference function.

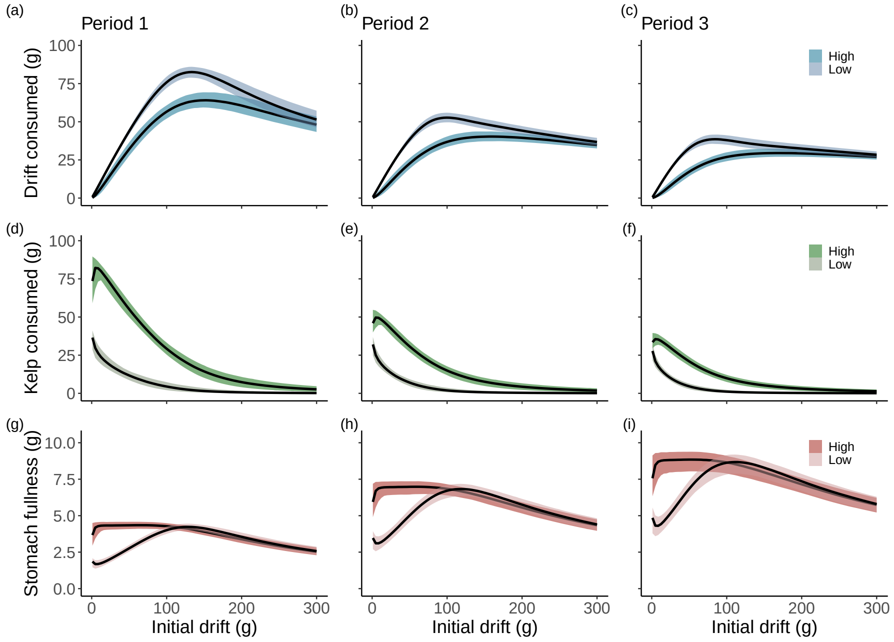

Figure S18: As in Fig. 3 of the main text for the logistic preference function with gut clearance ($z$).

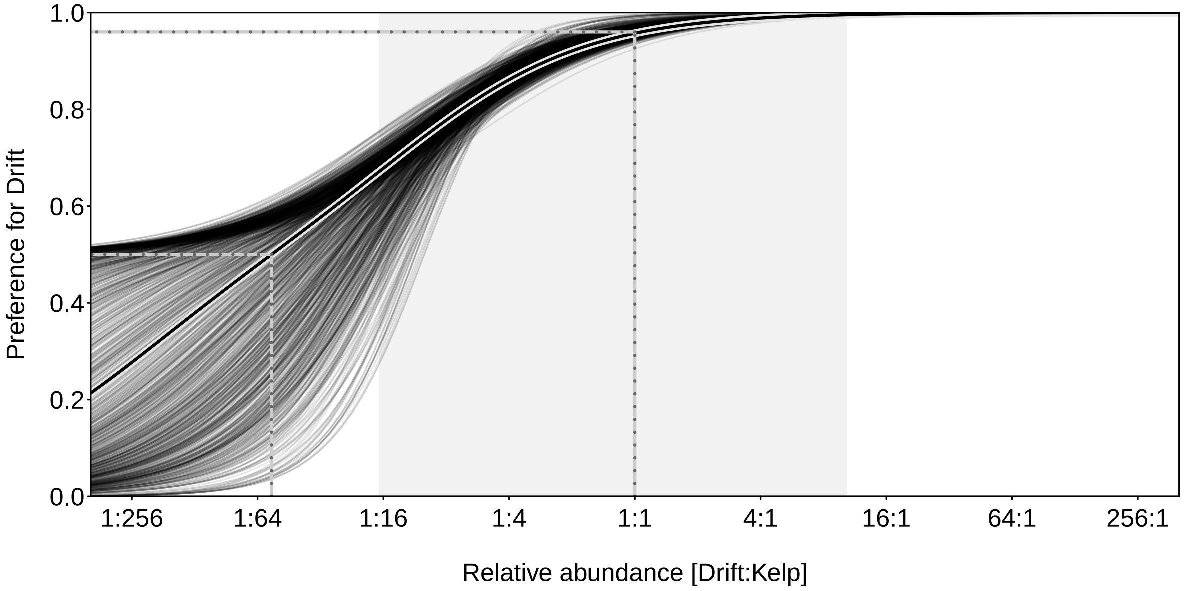

Figure S19: As for Fig. 4 of the main text for the reduced van Leeuwen preference function.

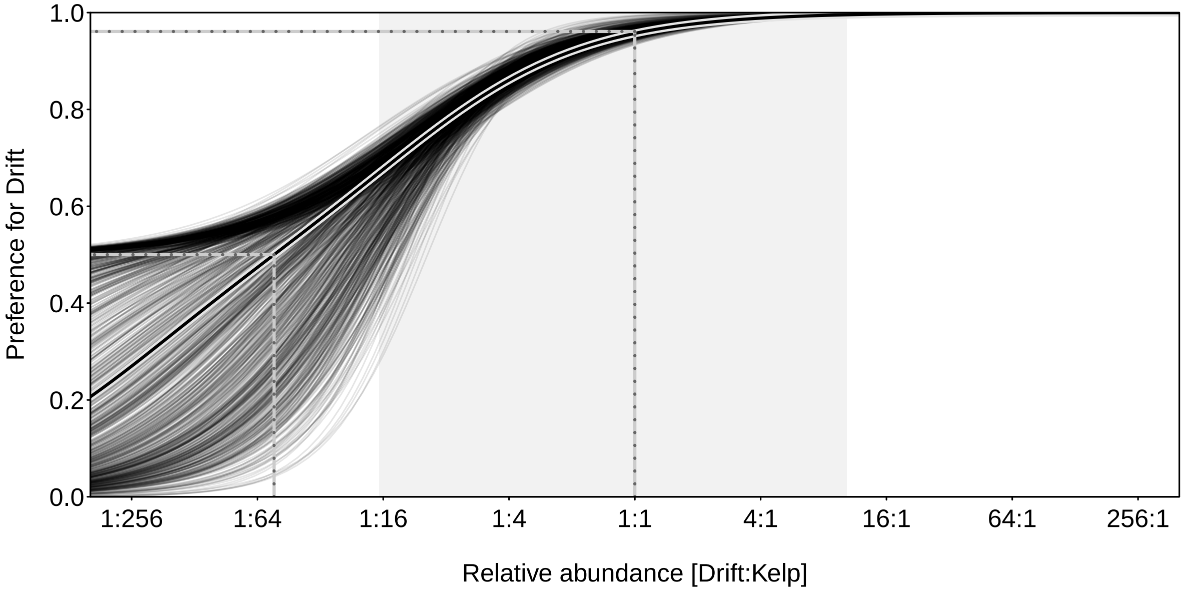

Figure S20: As for Fig. 4 of the main text for the reduced van Leeuwen preference function with gut clearance ($z$).

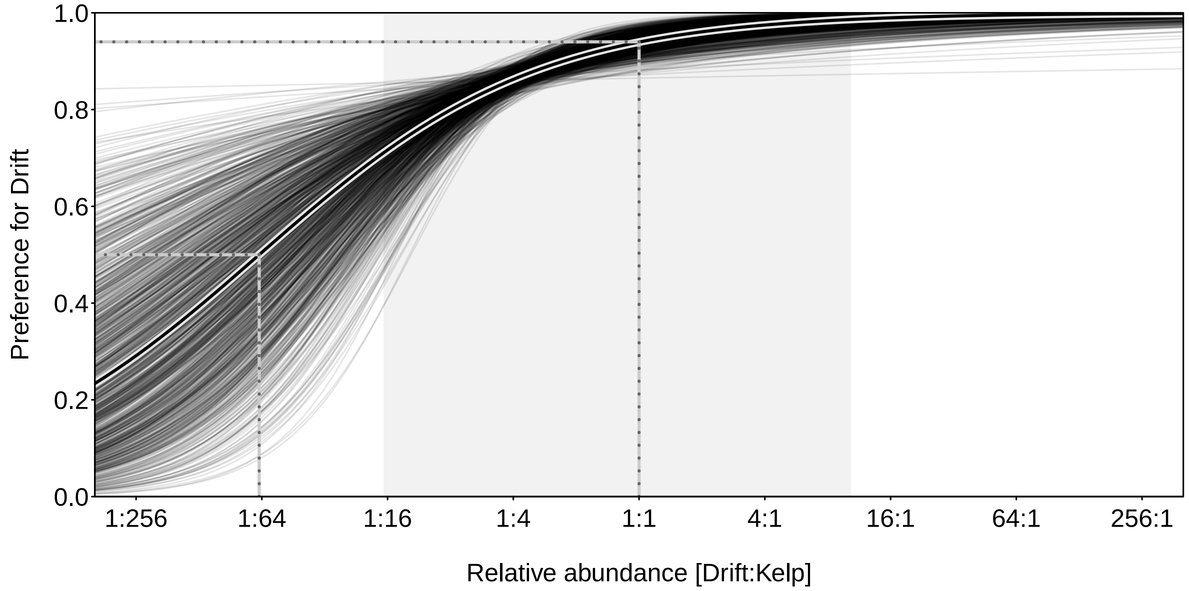

Figure S21: As for Fig. 4 of the main text for the logistic preference function.

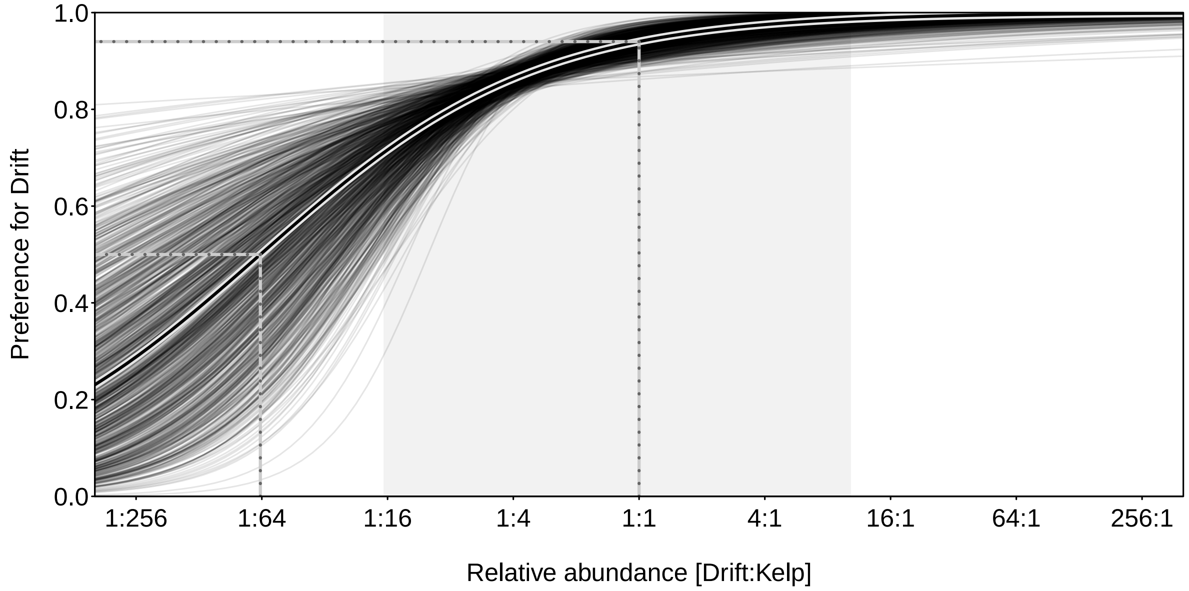

Figure S22: As for Fig. 4 of the main text for the logistic preference function with gut clearance ($z$).

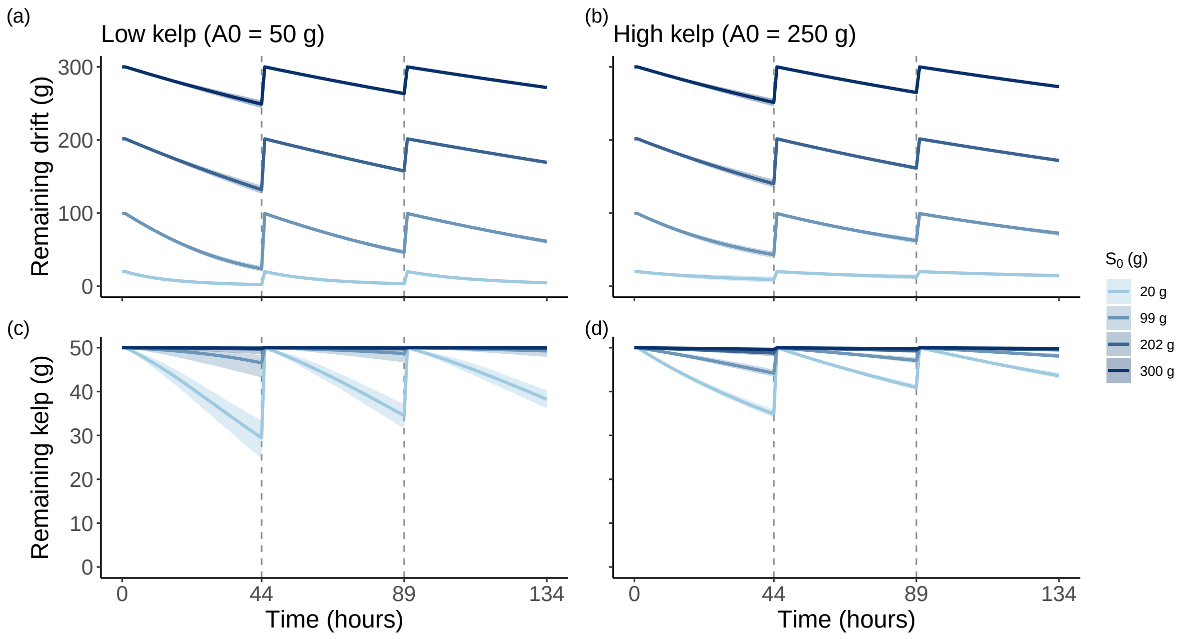

Figure S23: The model-average predicted temporal progression of remaining drift and kelp (with 95% credible intervals) for representative initial abundances of drift ($S_0$) and kelp ($A_0$).
